# Two distinct types of eye-head coupling in freely moving mice

**DOI:** 10.1101/2020.02.20.957712

**Authors:** Arne F. Meyer, John O’Keefe, Jasper Poort

## Abstract

Animals actively interact with their environment to gather sensory information. There is conflicting evidence about how mice use vision to sample their environment. During head restraint, mice make rapid eye movements strongly coupled between the eyes, similar to conjugate saccadic eye movements in humans. However, when mice are free to move their heads, eye movement patterns are more complex and often non-conjugate, with the eyes moving in opposite directions. Here, we combined eye tracking with head motion measurements in freely moving mice and found that both observations can be explained by the existence of two distinct types of coupling between eye and head movements. The first type comprised non-conjugate eye movements which systematically compensated for changes in head tilt to maintain approximately the same visual field relative to the horizontal ground plane. The second type of eye movements were conjugate and coupled to head yaw rotation to produce a “saccade and fixate” gaze pattern. During head initiated saccades, the eyes moved together in the same direction as the head, but during subsequent fixation moved in the opposite direction to the head to compensate for head rotation. This “saccade and fixate” pattern is similar to that seen in humans who use eye movements (with or without head movement) to rapidly shift gaze but in mice relies on combined eye and head movements. Indeed, the two types of eye movements very rarely occurred in the absence of head movements. Even in head-restrained mice, eye movements were invariably associated with attempted head motion. Both types of eye-head coupling were seen in freely moving mice during social interactions and a visually-guided object tracking task. Our results reveal that mice use a combination of head and eye movements to sample their environment and highlight the similarities and differences between eye movements in mice and humans.

**Highlights:**

- Tracking of eyes and head in freely moving mice reveals two types of eye-head coupling
- Eye/head tilt coupling aligns gaze to horizontal plane
- Rotational eye and head coupling produces a “saccade and fixate” gaze pattern with head leading the eye
- Both types of eye-head coupling are maintained during visually-guided behaviors
- Eye movements in head-restrained mice are related to attempted head movements

## Introduction

During natural behaviours, animals actively sample their sensory environment (Kleinfeld et al., 2006, Gottlieb and Oudeyer, 2018). For example, humans use a limited and highly structured set of head and eye movements (Sağlam et al., 2011, and references therein) to shift their gaze (gaze = eye-in-head + head-in-space) to selectively extract relevant information during visually-guided behaviors, like making a cup of tea (Land et al., 1999) or a peanut butter sandwich (Hayhoe, 2000). Revealing the precise patterns of these visual orienting behaviours is essential to understand the function of vision in humans and other animals (Hayhoe and Ballard, 2014, Land, 2015) and to investigate the underlying neural mechanisms.

The mouse has emerged as a major model organism in vision research, due to the availability of genetic tools to dissect neural circuits and model human disease. This has yielded detailed insights into the circuitry and response properties of early visual pathways in mice (see Seabrook et al. (2017) for a recent review). Mice use vision during natural behaviors such as threat detection (Yilmaz and Meister, 2013) and prey capture (Hoy et al., 2016). They can also be trained on standard visual paradigms similar to those used in humans and non-human primates including visual detection and discrimination tasks, with or without head restraint (Huberman and Niell, 2011, Carandini and Churchland, 2013, Horner et al., 2013). However, very little is known about how visual orienting behaviors support vision in mice. Vision in mice is typically studied in head-restrained animals to facilitate neural recordings and experimental control of visual input. Until recently, it has not been feasible to simultaneously measure movement of the head and eyes in freely behaving mice. The aim of our study was therefore to determine how eye and head movements contribute to visually-guided behaviors.

There is conflicting evidence about the role of eye movements in mice. Mice have laterally facing eyes with a large field of view of approximately 280° extending in front, above, below, and behind the animal’s head (Wagor et al., 1980, Dräger, 1978, Hübener, 2003, Seabrook et al., 2017, Samonds et al., 2018). There is only a narrow binocular field of approximately 40 − 50° overlap. In contrast to humans, mice have no fovea and appear to lack other pronounced retinal specializations for high resolution vision (Dräger and Olsen, 1981, Jeon et al., 1998). Despite this, multiple studies have found that head-restrained mice move their eyes (Sakatani and Isa, 2007, Wang et al., 2015, Samonds et al., 2018, Itokazu et al., 2018, Meyer et al., 2018); these eye movements are rapid and conjugate, i.e. both eyes moving together in the same direction, with an average magnitude of 10 − 20° and peak velocities that can reach more than 1000°/s. While these saccade-like eye movements provide only a relatively small shift in the visual field (about 5 %), mainly in the horizontal direction (Sakatani and Isa, 2007), it has been suggested that they resemble exploratory saccades in humans (Sakatani and Isa, 2007, Samonds et al., 2018).

However, studies in freely moving mice (Payne and Raymond, 2017, Meyer et al., 2018) and also rats (Wallace et al., 2013) have found eye movement patterns much more complex and often non-conjugate, i.e. both eyes moving in opposite directions. These non-conjugate eye movements were systematically linked to changes in orientation of the animal’s head with respect to the horizontal plane (head tilt) (Oommen and Stahl, 2008, Wallace et al., 2013, Meyer et al., 2018). While the precise function of this eye-head coupling is still unclear (but see Wallace et al. (2013), Meister and Cox (2013)), it appears to be largely compensatory and has been suggested to serve to stabilize the visual field with respect to the ground (Oommen and Stahl, 2008, Meyer et al., 2018).

We report that saccades and head orientation-related changes in eye position simultaneously serve two distinct and complementary functions in freely moving mice. We previously observed that freely moving mice rarely make saccades in the absence of head motion (Meyer et al., 2018). We therefore reasoned that saccades might serve to shift the gaze during combined eye and head movement, similar to higher vertebrates including humans, primates, cats and rabbits (Land, 2019). At the same time, compensation for changes in head orientation could then approximately maintain the same view of the visual environment with respect to the horizontal ground plane, consistent with previous observations that head orientation accounts for most variability in the vertical eye axis, but fails to account for a substantial fraction of variability in the horizontal eye axis along which saccadic eye movements mainly occur in mice (Meyer et al., 2018).

To investigate eye/head movement relations we used a system that we recently developed for tracking eye positions together with head tilt and head rotations in freely moving mice (Meyer et al., 2018). We show that eye movements can be decomposed into non-conjugate head tilt-related and conjugate eye movements along the horizontal eye axis. Non-conjugate changes in eye position occur during head tilt and stabilize the gaze of the two eyes relative to the horizontal plane. In contrast, conjugate horizontal eye movements yielded a “saccade and fixate” gaze pattern that was closely linked to rotational head movements around the yaw axis. Eye movements during the “saccade” and the “fixate” phases were strongly coupled to the head but in different rotation directions and this coupling was preserved when animals were engaged in a novel visually-guided tracking task. Indeed, eye movements in head-restrained mice always occurred during attempted head movements, and the direction of the attempted head movement was consistent with that of combined eye-head gaze shifts in freely moving animals.

Our results resolve the apparent discrepancy between eye movement patterns in head-restrained and freely moving mice. To summarize, eye movements in mice consist of two distinct, separable types: non-conjugate head tilt-related and conjugate eye movements along the horizontal eye axis. Importantly, gaze shifts in mice rely on combined head and eye movements with a similar “saccade and fixate” pattern as in other higher vertebrates, including humans.

## Results

### Eye Movements in Freely Moving Mice, Head-restrained Mice, and Humans

To investigate how mice use their head and eyes to explore the environment we tracked the positions of both eyes together with head motion in freely moving mice using a previously developed head-mounted system (Meyer et al., 2018). The system includes two head-mounted cameras combined with an inertial measurement unit (IMU) sensor (Figure 1A). The IMU provides information about head tilt and head rotation while the cameras measure the positions of the eyes relative to the eye axis in the head coordinate frame.

**Figure 1:**
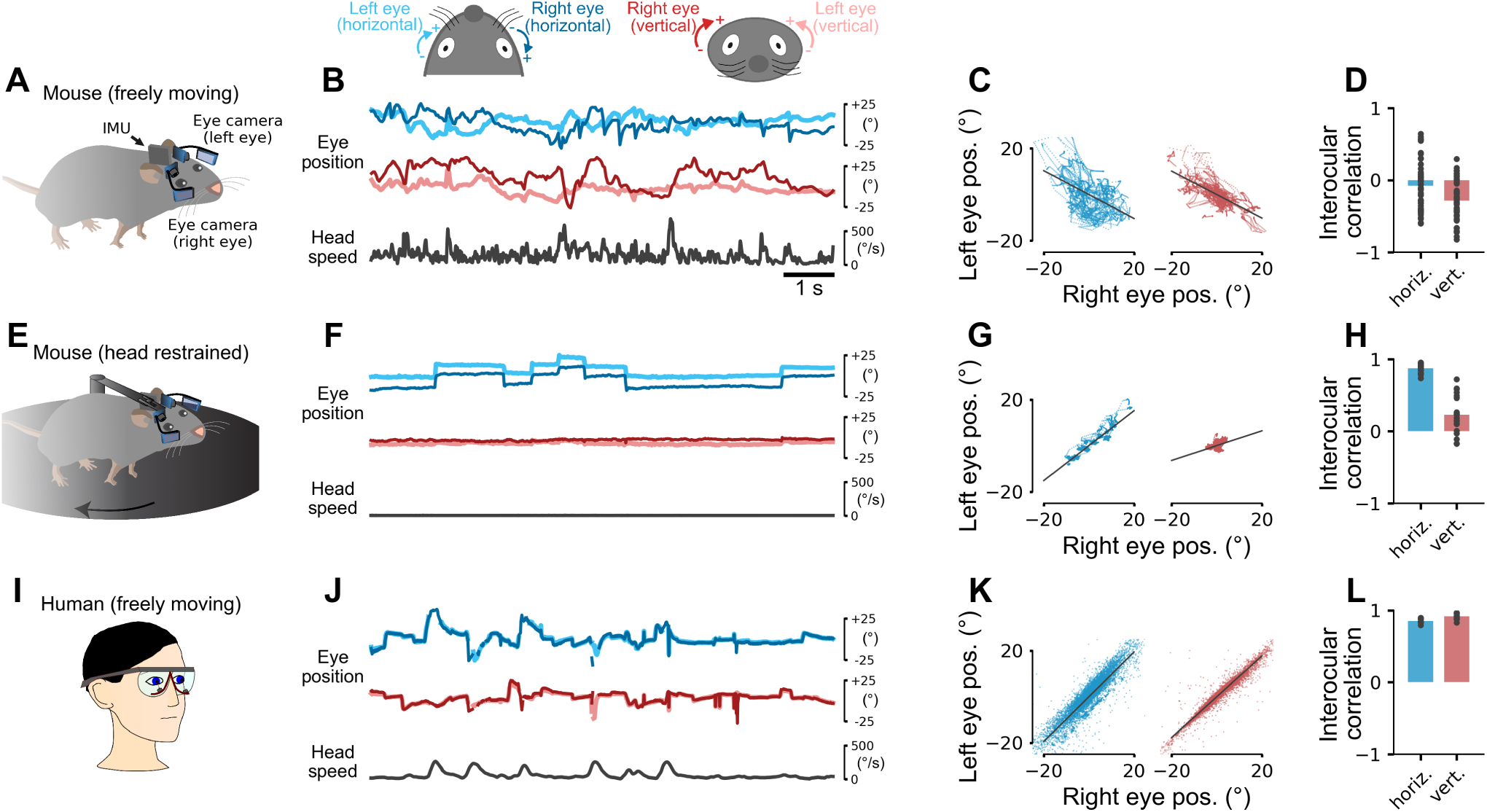
Eye Movements in Freely Moving Mice, Head-restrained Mice, and Humans. (A) Tracking eye and head motion in a freely moving mouse. Videos of each eye are recorded using miniature cameras and infrared (IR) mirrors mounted on an implant with a custom holder. Each eye is illuminated by two IR light sources attached to the holder. The mirrors reflect only IR light and allow visible light to pass so that the animal’s vision is not obstructed. Head motion and orientation are measured using an inertial measurement unit (IMU). (B) Eye coordinate systems used in this study (top). A 10 second example segment showing horizontal and vertical position of both eyes and head speed in an unrestrained, spontaneously behaving mouse (bottom). (C) Horizontal (left) and vertical (right) eye positions for the whole recording of the data in B (10 minutes). Interocular eye positions were negatively correlated (solid black line) and on average (D) small and negative. (E) Eye tracking in a head-restrained mouse on a running disk using the same technique as in A. (F-H) The same as in B–D but for a head-restrained mouse. In contrast to the freely moving condition, eye movements mostly occurred in the horizontal direction and were tightly coupled between the eyes. (I) Tracking eye and head movement in freely moving humans, using goggles with integrated eye tracking cameras and IR illumination. (J–L) The same as in B-D but for humans walking through the environment. Interocular correlations between the two eyes in humans show strong coupling between horizontal and vertical eye positions. Note that the lines in J for left and right eye positions are closely overlapping. Time scale in F and J same as B. See also Video S1.

We defined the horizontal eye coordinate system in mice and humans with clockwise positions more positive in each eye (Figure 1B, top). For the left eye, horizontal eye positions closer to the nose have more positive values while for the right eye, horizontal eye positions further away from the nose are more positive. In the vertical direction, eye positions of both eyes further towards the top of the eye are more positive. With this coordinate system, conjugate eye movements (typical in humans) generate positive correlations between horizontal and vertical eye positions of the two eyes, while non-conjugate eye movements generate negative ones.

Eye movements in mice freely exploring an environment showed large horizontal and vertical displacements of the two eyes (Figure 1B, bottom). On average, these displacements were weakly correlated across the two eyes (Figure 1C,D; *r* = −0.07 ± 0.05 horizontal, *r* = −0.29 ± 0.04 vertical, *n* = 47 recordings in 5 mice, 10 min each). In contrast, when the same mice were head-restrained (i.e. the head was fixed but the mice free to run on a wheel, Figure 1E) both eyes showed saccadic-like eye movements, preferentially in the horizontal direction (Figure 1F) with high interocular correlations (Figure 1G,H; *r* = 0.88 ± 0.01 horizontal and *r* = 0.23 ± 0.06 vertical eye positions; *n* = 20 recordings from 2 mice, 10 min each). Thus, eye movement patterns differed substantially in freely moving and head-restrained mice.

For comparison, we also recorded eye movements in humans walking around an environment using commercially-available head-mounted eye tracking goggles (Figure 1I). Eye positions were strongly correlated between both eyes (Figure 1J–L; interocular correlations *r* = 0.85 ± 0.01 horizontal and *r* = 0.92 ± 0.01 vertical, *n* = 10 recordings from 5 subjects, recording time 427 ± 200 s). Thus, eye movements in freely moving mice differed substantially from eye movements in humans. In contrast, head-restrained mice made saccadic-like eye movements strongly coupled across the two eyes, similar to those in humans. An obvious difference between the two conditions in the mouse was that they moved their heads a lot during free exploration which was not possible during head restraint (Figure 1B,F). Coupling of the eyes to the motion of the head, therefore, could be a potential explanation for the observed differences.

### Head Tilt-related Changes in Eye Position Stabilize Gaze Relative to the Horizontal Plane

We first analyzed the effect of head tilt on eye position in freely moving mice. Previous results in head-restrained (Andreescu et al., 2005, Oommen and Stahl, 2008) and freely moving mice (Meyer et al., 2018) had suggested that average eye position systematically varies with the tilt of the head (combined pitch and roll). To test if this were also true for our data, we computed average eye position separately during either pitch or roll of the head (Figure 2A–D). Head pitch had an effect on both horizontal and vertical eye position (Figure 2A,B) whereas roll predominantly affected vertical eye position (Figure 2C,D). During upward (positive) head pitch, both eyes turn downwards and inwards towards the nose (Figure 2B, top). In contrast, during downward (negative) head pitch, the opposite happens, both eyes turn upwards and outwards towards the ears (Figure 2B, bottom). During positive roll (lowering the right side of the head relative to the left side), the right eye moves upward and the left eye moves downward (Figure 2D, top); the opposite happens during negative head roll (Figure 2D, bottom).

**Figure 2:**
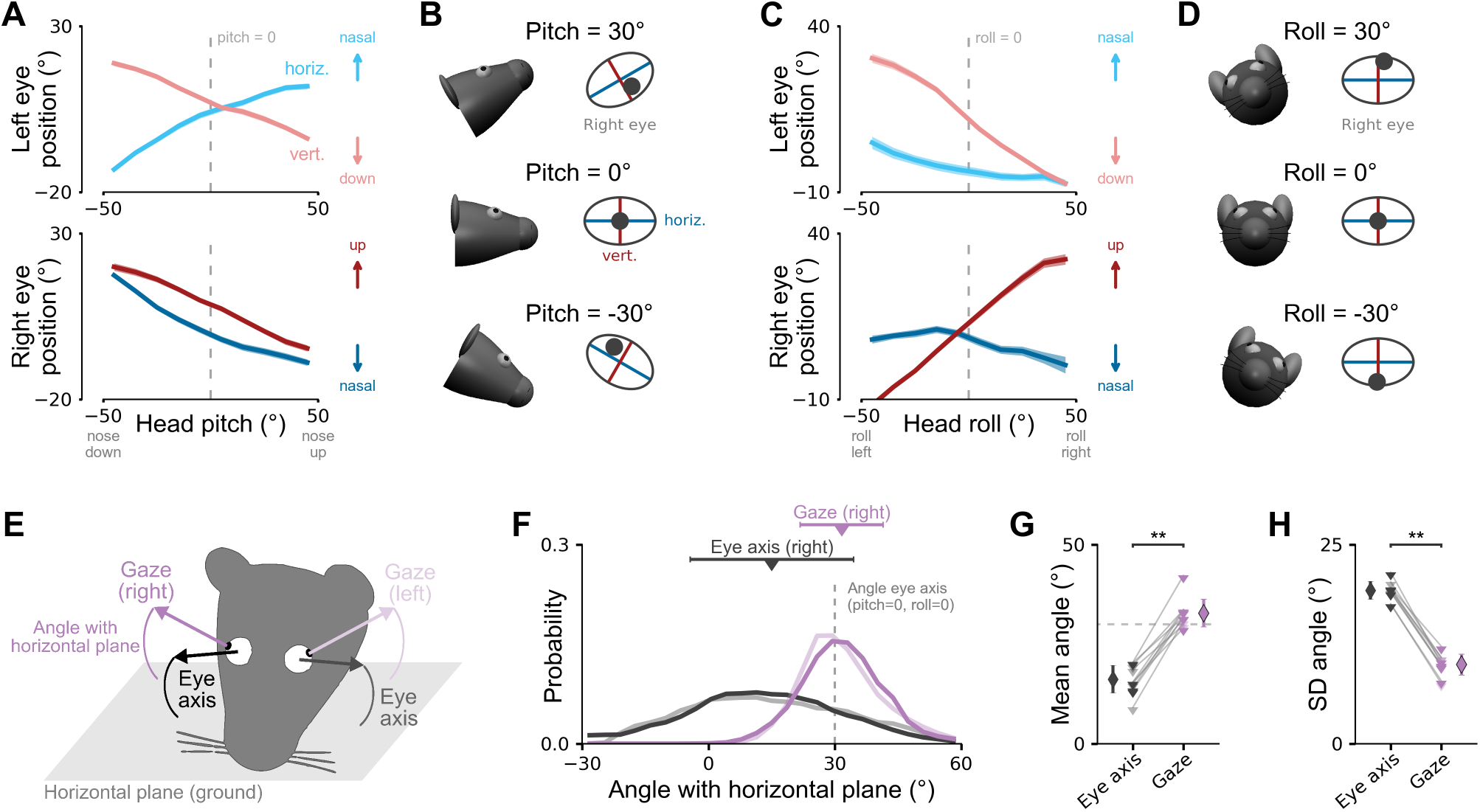
Head Tilt-related Changes in Eye Position Stabilize Gaze Relative to the Horizontal Plane. (A) Horizontal (blue lines) and vertical eye position (red lines) as a function of head pitch for the left (top) and right eye (bottom). Arrows indicate directions of eye position change in the eye coordinate system. Dashed vertical line shows pitch = 0°. (B) Illustration of systematic dependence of horizontal and vertical eye position on head pitch for different pitch values. For illustration, intersection of horizontal and vertical eye axes aligned with average eye position for pitch = 0° (dashed line in A). (C and D) The same as in A and B but as a function of head roll. Same data as in Figure 1D. (E) Illustration of eye axes fixed in a head-centered reference frame (black arrow) and gaze axes (center of pupil rotating in head; violet arrow) for left and right eyes. Angles of axes relative to horizontal plane (ground; gray area). (F) Distributions of angles of eye axes (black/gray lines) and gaze axes (violet lines) with horizontal plane for one example mouse. Negative angles indicate axis pointing downwards (to the horizontal plane) whereas positive angles indicate upward pointing axis. For reference, angle of eye axis for pitch = 0° and roll = 0° is shown (dashed gray line). Triangles and bars indicate circular mean and standard deviation of distributions, respectively. Same color scheme as in E. (G) Circular mean angles for left and right eye in 5 mice. Same color scheme as in E. (H) Circular standard deviation of angles for the same data. Diamonds represent mean and standard deviation across mice (left and right eye). Same data as in A and C. See also Figure S1 and Video S2.

The observed effect of head tilt on eye position is consistent with a stabilization scheme (Oommen and Stahl, 2008, Meyer et al., 2018) that influences the position of the eyes with regard to gravity (based on vestibular input) (Andreescu et al., 2005, Oommen and Stahl, 2008). We therefore reasoned that one potential function of changes in eye position could be alignment of the visual field with the horizontal plane (ground). To directly test this, we calculated the angle between the “gaze”, i.e. the vector determined by the center of the pupil rotating in the head, and the horizontal plane (Figure 2E). The calculation of gaze angles involved a geometric model of the eye axes in the head because of misalignment of the eye and the head (pitch and roll) axes (Figure S1A and STAR Methods). Gaze angles for both eyes were typically positive (i.e. pointing slightly upwards from the horizontal plane) and tightly centered around the angle of the eye axis when the animal kept its head straight (pitch = 0° and roll = 0°; Figure 2F–H; gaze angle mean = 32.8 ± 3.5°, SD = 9.9 ± 1.3°). In contrast, the angle of the eye axis, defined as the origin of the eye coordinate system fixed in a head-centered reference frame (Figure 2B,D), showed a much wider distribution with the axis frequently pointing towards the ground (Figure 2F–H; eye axis angle mean = 16.2 ± 3.5°, SD = 19.3 ± 1.1°; *p* = 0.002, gaze vs eye axis mean; *p* = 0.002, gaze vs eye axis SD; Wilcoxon signed-rank tests, *n* = 10 (5 mice, left and right eye)). This suggests that one function of this head tilt-related eye-head coupling in the mouse could be stabilization of the visual field relative to the horizontal plane.

### Horizontal Eye Movements not Explained by Head Tilt Are Conjugate across the Two Eyes

Next, we investigated whether eye movements that were not explained by head tilt revealed some properties of the saccadic-like eye movements observed in head-restrained mice. To isolate the head tilt-related component, and to reveal the component not explained by head tilt, we took advantage of the accelerometer signals of the head-mounted IMU to measure head pitch and roll, and used these to predict horizontal and vertical eye positions for each eye using regression models (Figure 3A; see also Meyer et al. (2018)). Most of the variance in eye position could be explained by head pitch and roll (Figure 3B). Model predictions were significantly more accurate for vertical than horizontal eye positions (*r*^2^ = 0.86 ± 0.01 vertical, *r*^2^ = 0.62 ± 0.02 horizontal, *p* = 1 · 10^−16^, Wilcoxon signed-rank test; *n* = 47 recordings in 5 animals, 10 min each).

**Figure 3:**
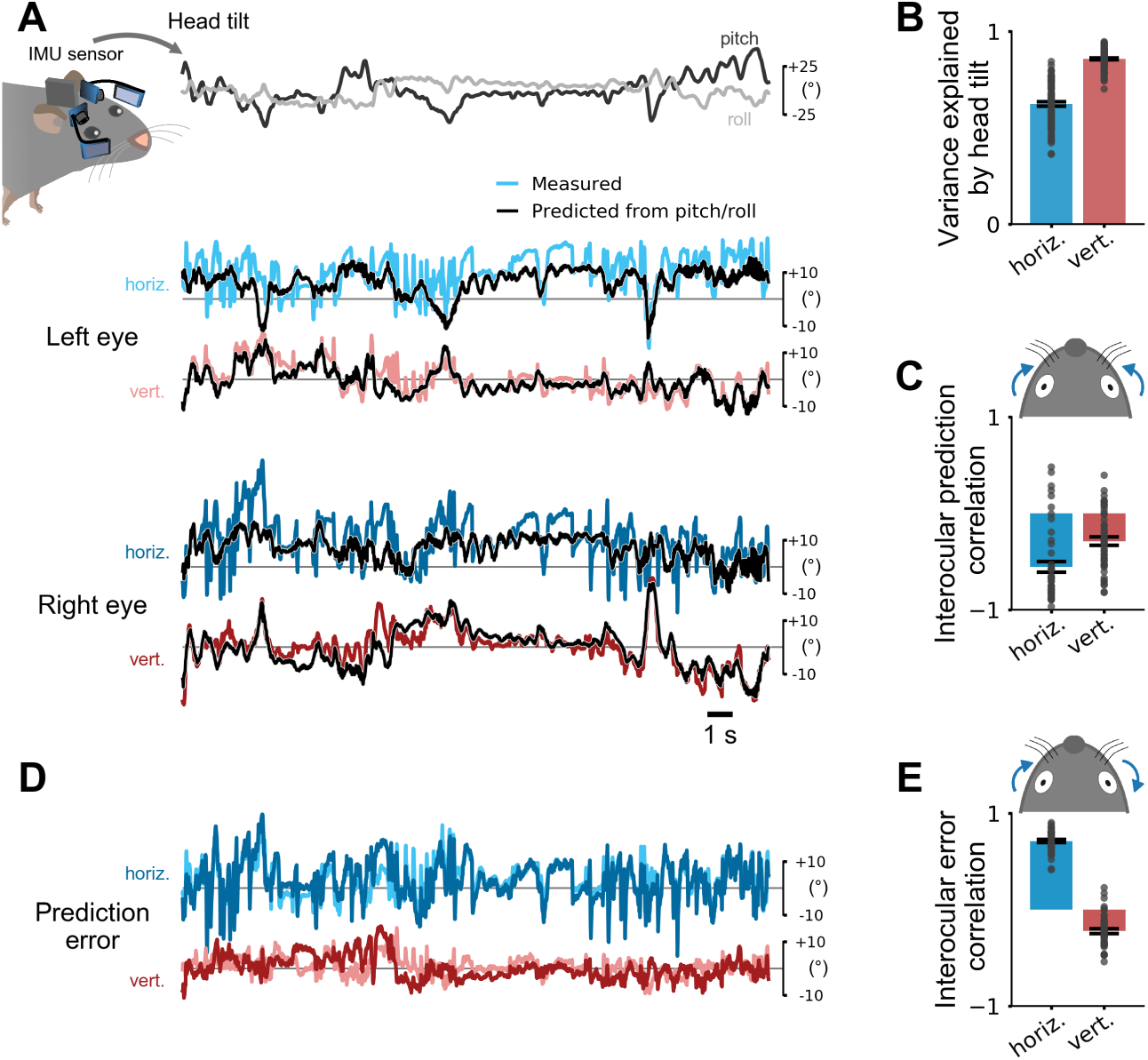
Horizontal Eye Movements not Explained by Head Tilt Are Conjugate across the Two Eyes. (A) Top: Head tilt was measured using the IMU sensor attached to the animal’s head. Eye positions were measured using the head-mounted camera system. Computational models were used to predict horizontal and vertical eye positions from head pitch and roll for each eye. Bottom: Measured (colored lines) and predicted (black lines) horizontal and vertical eye positions for both eyes. (B) Cross-validated explained variance along the horizontal (horiz.) and vertical (vert.) eye axes (*n* = 47 recordings from 5 mice, 10 min each). Head tilt explained 86% variance in vertical but only 62% in horizontal eye position. Recordings for each eye axis pooled across eyes and mice. (C) Interocular correlation of the eye movements that were predictable by head pitch and roll (i.e. the predictions of independent models for the two eyes as shown in A). Strong negative correlation for horizontal eye movements indicates convergence and divergence across eyes. Blue arrows show horizontal convergence. Same data as in B. (D) Prediction errors for the eye position traces in A showed strong co-fluctuations in horizontal but not vertical eye direction. (E) Interocular correlation of the eye movements that were not predictable by head pitch and roll (i.e. the prediction errors of independent models for the two eyes as shown in D). There was a strong positive correlation for horizontal eye movements suggesting that conjugate eye movements occurred during head free behavior and were not explained by head tilt. Arrows show coupling for left eye rotating in nasal direction. Same data as in B.

We wondered if the lower predictability of horizontal eye position by head tilt might be due to the inclusion of conjugate eye movements similar to those observed in head-restrained mice. Changes in head pitch have been shown to be associated with convergent horizontal eye movements, i.e. both eyes rotate towards the nasal edge when the head pitches up, and divergent eye movements, i.e. both eyes rotate towards the temporal edge when the head pitches down (Oommen and Stahl, 2008, Wallace et al., 2013, Meyer et al., 2018). As a consequence, pupil position will be negatively correlated across the two eyes. In contrast, conjugate eye movements, such as the ones observed in head-restrained mice, will result in positive interocular correlations; both eyes rotate either clockwise (CW) or counter-clockwise (CCW), e.g., one eye moves towards the nasal edge while one moves away from the nasal edge. We therefore reasoned that, if conjugate eye movements occur in freely moving mice, failure to predict eye position based on head tilt should lead to positive rather than negative interocular correlations.

Indeed, consideration of head tilt succeeded in separating two different types of horizontal eye movement: positions predicted by models based on head pitch and roll were consistent with convergent/divergent horizontal eye movements, i.e. both eyes tended to move together towards the nasal or temporal edge (interocular correlation *r* = −0.56 ± 0.06; Figure 3C); those not predicted by head tilt (prediction error, Figure 3D) were positively correlated, implying conjugate horizontal eye movements (interocular correlation *r* = 0.71 ± 0.02; Figure 3E). Thus, conjugate horizontal eye movements, potentially resembling those observed in head-restrained mice, are also part of the natural repertoire in freely moving mice and co-occur with non-conjugate eye movements.

### Rapid Saccadic Conjugate Horizontal Eye Movements in Freely Moving Mice

We next identified rapid saccadic-like horizontal eye movements in freely moving mice similar to those in head-restrained mice (Figure 1E–H). Viewed on a fast timescale, the two eyes often showed brief, strong co-fluctuations (Figure 4A, top) of high velocities frequently reaching more than 800 °/s (Figure 4A, middle). These saccadic eye movements were not only strongly coupled across the two eyes but also conjugate (Figure 4B), similar to saccades in head-restrained animals. In total, during 94% (9703/10331) of all saccades detected for both eyes, the two eyes were moving in the same direction (CW or CCW). Saccades in freely moving mice, therefore, were qualitatively similar to those in head-restrained mice.

**Figure 4:**
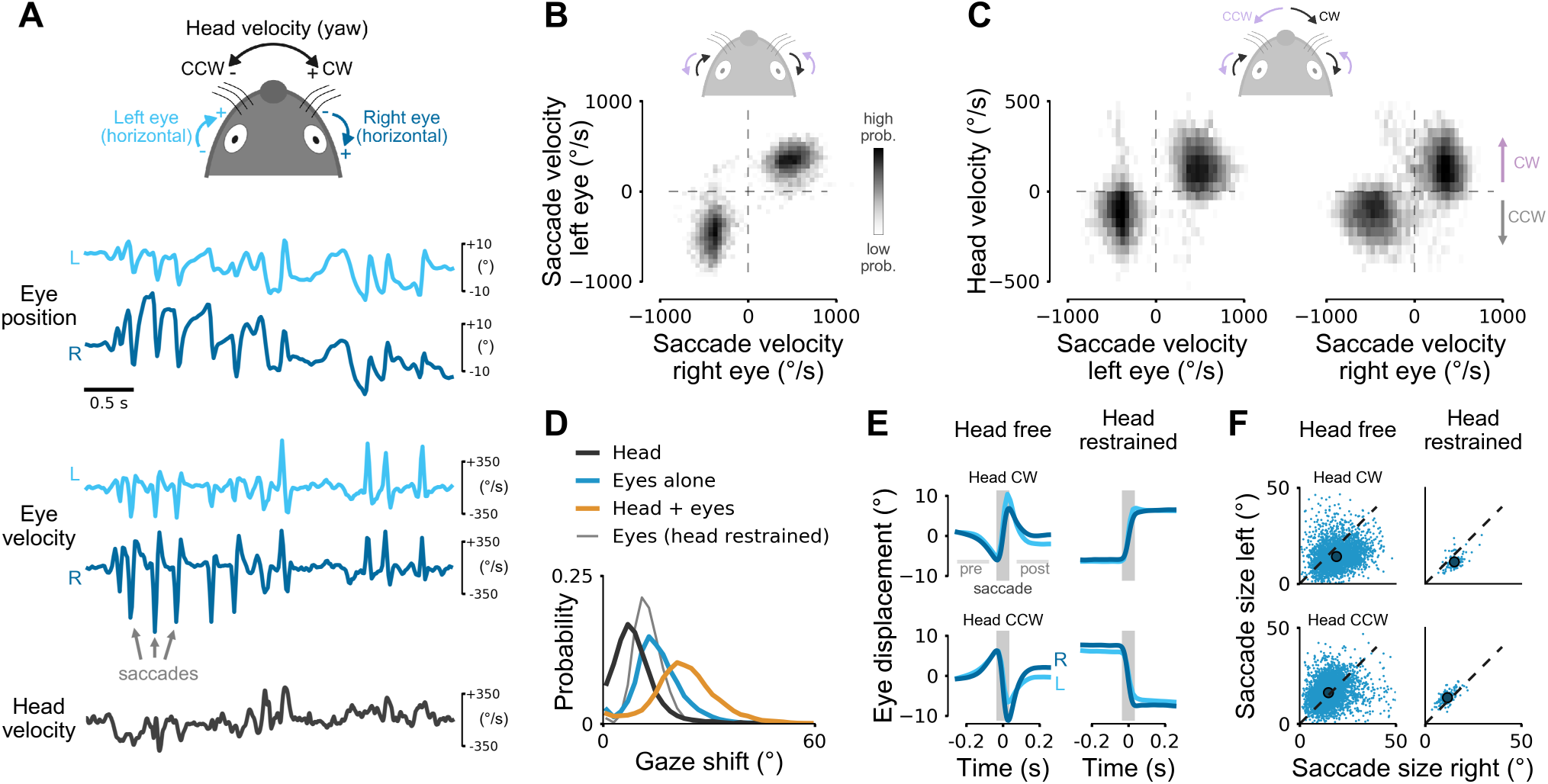
Rapid Saccadic Conjugate Horizontal Eye Movements in Freely Moving Mice. (A) Schema for representing horizontal head and eye rotation axes (top). Examples of eye position (top traces), angular eye velocity (middle), and angular head velocity (bottom) in a freely moving mouse. High-velocity peaks during saccades coordinated between the two eyes. Left eye in light blue, right eye in dark blue. (B) Log-scaled joint distribution of horizontal saccade velocity for the left and right eyes. 94 % of saccades had the same sign for both eyes (10331 saccades detected in both eyes). (C) Log-scaled joint distribution of horizontal saccade velocity for the left and right eyes and angular head velocity. Most saccades occurred during head rotations with eye rotations in the same direction as the head, same data as in Figure 1B. (D) Gaze shift magnitudes during saccades in freely moving mice, eyes and head together in orange, eyes alone in blue, head alone in dark gray, and for comparison, gaze shift in head-restrained mice, thin gray line. (E) Average saccade profiles in freely moving (left) and head-restrained (right) mice. Saccades for clockwise (CW, top) and counter-clockwise (CCW, bottom) head rotations (shaded gray area) were preceded and followed by a counter eye movement (“pre” and “post”) in freely moving mice but not in head-restrained mice. Means ± SEM (smaller than line width). (F) Saccade sizes for CW and CCW head rotations in head free (left) mice. On average, saccades were larger for temporal-to-nasal than for nasal-to-temporal saccades. Dots indicate average saccade sizes. Saccades in head-restrained mice for comparison (right) with same asymmetry in average saccade sizes as in head free mice.

In many species, including humans (Guitton and Volle, 1987), cats (Guitton et al., 1984, Guitton, 1992), and rabbits (Collewijn, 1977), horizontal eye movements are often linked to rotations of the head. We therefore tested whether this is also true in mice. During saccades, eyes and head rotated in the same direction (i.e. CW or CCW, Figure 4C) similar to the pattern observed in freely moving humans (Figure S2A,B). We compared the effect of combined eye and head movements to eye or head shifts alone (integrated head yaw velocity signal from the head-mounted gyroscope). We found that eye or head movements alone shifted gaze by 15.8 ± 7.0 ° and 9.4 ± 6.5 °, respectively (Figure 4D). Head motion thus substantially contributed to gaze shifts which in total equalled 23.3 ± 9.6 °. Saccades in head-restrained mice were considerably smaller (12.7 ± 4.0 °, 460 saccades from 2 mice) than gaze shifts or even eye saccades alone in head free mice (*p* = 6.6 · 10^−16^, “Head + eyes” vs “Eyes (head restrained)”; *p* = 2.4 · 10^−16^, “Eyes alone” vs “Eyes (head restrained)”; Wilcoxon rank-sum test, Bonferroni correction).

Gaze shifts in many animals are often observed together with periods of gaze stabilization (Land, 2015), and we wondered if a similar pattern could be observed in mice. To test this, we computed average displacements of both eyes, aligned to saccade onset (Figure 4E, left). These traces revealed two distinct features. First, saccades were preceded and followed by slower counter movements of the eyes (as indicated by eye displacement in opposite direction to the saccade in Figure 4E, left). This suggests that eye movements in freely moving mice not only support gaze shifts but also help to stabilize the image of the surrounding just before and after saccades during head rotation. The absence of counter movements in head-restrained mice (Figure 4E, right) further supports this hypothesis. Second, while the temporal profile of saccades in both eyes was similar, saccade size was larger for temporal-to-nasal than for nasal-to-temporal movement (Figure 4E,F, left; *p* = 1 · 10^−16^, CW left vs. right eye; *p* = 1 · 10^−16^, CCW left vs. right eye; Wilcoxon signed-rank tests). Consistent with previous studies (Sakatani and Isa, 2007, Itokazu et al., 2018), we found a similar pattern for head-restrained mice (Figure 4E,F, right; *p* = 5.4 · 10^−14^, CW left vs. right eye; *p* = 1.4 · 10^−15^, CCW left vs. right eye; Wilcoxon signed-rank tests) suggesting that saccades in head-restrained and freely moving mice share similar mechanisms. At the same time, this degree of asymmetry in saccade sizes and velocities between both eyes represents a major difference between mice and humans (Collewijn et al., 1988).

### Head and Eyes Contribute to a “Saccade and Fixate” Gaze Pattern

Gaze in mice involves “saccade and fixate” periods during which eye and head rotate together (gaze shift) preceded and followed by gaze stabilization (during which the eyes counter-rotate). We further investigated these patterns during natural behavior by comparing horizontal angular head position computed by integrating yaw velocity to horizontal eye positions using the head-mounted cameras.

Head position varied smoothly with large excursions of several hundred degrees in CW and CCW directions (Figure 5A, middle). In contrast, eye positions appeared jerky with both saccadic and slower movements of smaller amplitudes compared to the head rotations (Figure 5A, bottom). Combining head and eye positions to compute gaze (eye-in-head + head-in-space) revealed a step-like gaze pattern consisting of gaze shifts and periods during which the image of the external world was approximately stable (Figure 5A, top; Figure S3A,B). This pattern implies that mice view their surrounding as a sequence of stable images interrupted by rapid gaze shifts (1–2 shifts per second; Figure S3), similar to the stable view in humans that is only briefly interrupted by rapid saccadic eye movements.

**Figure 5:**
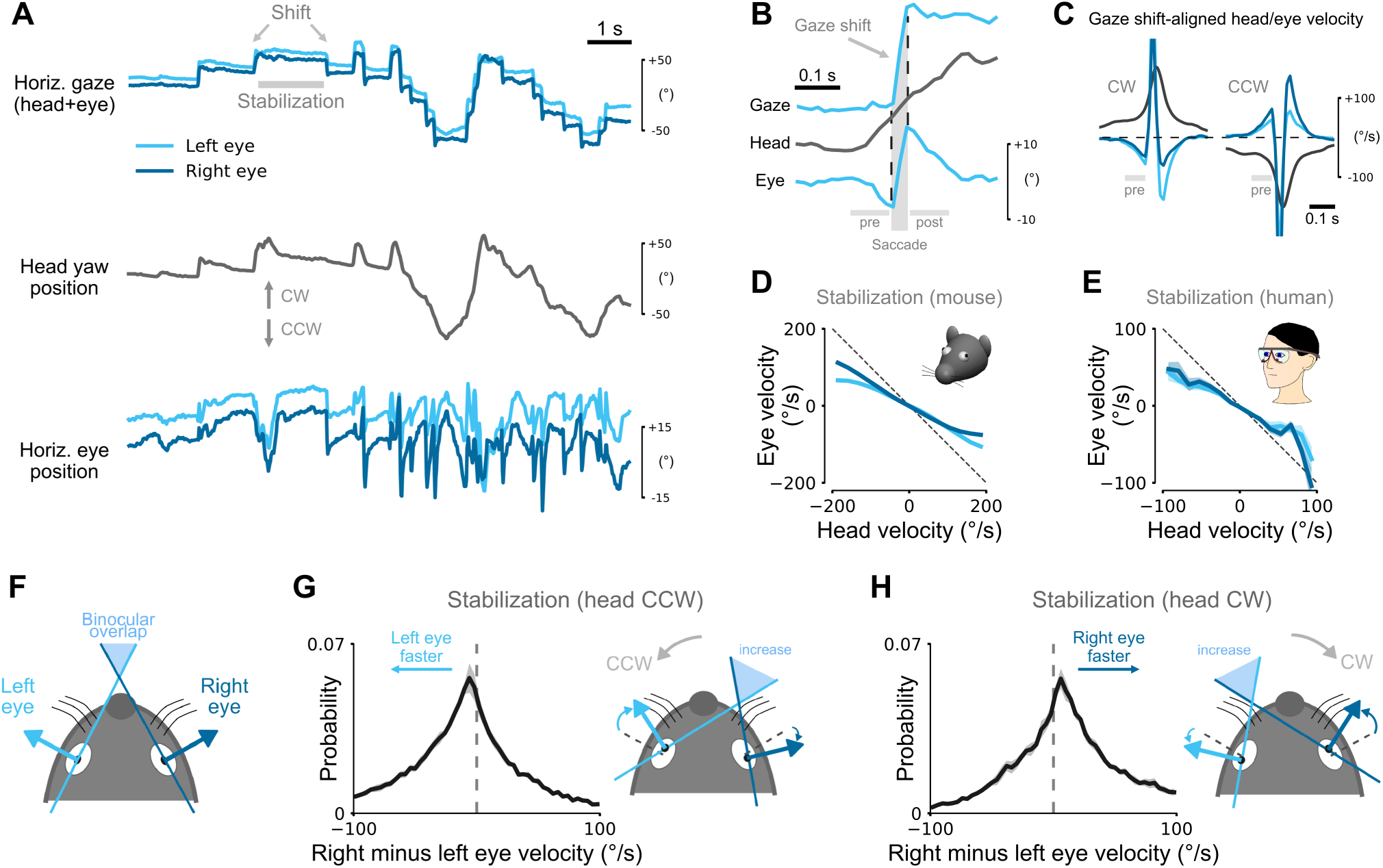
Head and Eyes Contribute to a “Saccade and Fixate” Gaze Pattern. (A) Horizontal positions of the two eyes (bottom), angular head yaw position (middle), and gaze (head + eye, top) during 12 s segment in a freely moving mouse selected to highlight the “saccade and fixate” pattern. Small amplitude, jerky eye movements and large amplitude, smooth head movements combine to produce the “saccade and fixate” gaze pattern. (B) Magnified traces for a single gaze shift from the recording in A. Head movement is accompanied by an initial counter-rotation of the eye before the gaze saccade. Vertical and horizontal gray bars indicate saccade and pre/post periods, respectively. (C) Gaze shift-aligned head and eye velocity traces for clockwise (CW, left) and counter-clockwise (CCW, right) gaze shifts. 44396 gaze shifts from 5 mice (22226 CW, 22170 CCW), mean ± SEM (smaller than line width). (D) Relation between horizontal eye and head velocity during stabilization periods (example period marked in A). Eye movements between saccadic gaze shifts counteract head rotations, mean ± SEM (smaller than line width). Dashed line indicates complete offsetting counter-rotation; same data as in Figure 1C. (E) The same as in B but for humans wearing head-mounted eye-goggles. (F) Illustration of monocular left and right visual field (about 180°) and horizontal binocular overlap. (G) Left: distribution of the difference in right and left eye velocity during stabilizing eye movements (CCW head rotation), mean ± SEM for 5 mice. Right: illustration of consequence of asymmetric nasal-to-temporal and temporal-to-nasal eye velocity on binocular overlap (increase relative to setting shown in F). (H) The same as in G but for CW head rotations.

We looked more closely at the precise sequence of head and eye movements during “saccade and fixate” gaze shifts. Figure 5B shows a close up of an example gaze shift. Consistent with the average saccade traces (Figure 4E), there were three distinct phases of eye motion: an initial eye movement that counter-acted the start of the head movement; the saccadic eye movement that is responsible for the rapid onset of the gaze shift; and a final “compensatory” phase, in which the eye rotates in a direction approximately equal and opposite to head rotation in order to ensure gaze stability. This three phase pattern was consistent across a large number of saccade and fixate shifts for a total of 22226 CW and 22170 CCW gaze shifts (Figure 5C). This uniformity suggests that head, and not eye, movement is the driver of gaze shifts in freely moving mice.

We turned our attention to the conjugate movements of the eyes during the “fixate” phase and asked to what extent they counteracted head yaw rotations to stabilize retinal images. To investigate this we compared angular velocities of eye and head in between the gaze shifts. Eyes were typically moving in the direction opposite to the head (Figure 5D), consistent with the expected effects of the angular vestibulo-ocular reflex (VOR). This relation was approximately linear and we quantified the mean absolute deviation (MAD) from full counter-rotation (dashed line in Figure 5D) and the “gain”, defined here as the negative slope of a line fitted to the data using linear regression. Across the measured velocity range, the MAD was 41.46 ± 7.82°/s with a gain of 0.53 ± 0.06. We repeated the same analysis for our human eye tracking data and found remarkably similar values in humans (Figure 5E; MAD 25.29 ± 4.74°/s, gain 0.59 ± 0.16); mice have a slightly reduced degree of image stabilization compared to humans. Changes in head tilt (Figure 3) had only a small and statistically insignificant impact on these parameters (Figure S3C–E), suggesting that head tilt-related visual field stabilization and gaze stabilization during the “fixate” phase of the “saccade and fixate” pattern act largely independent of each other in freely moving mice.

Finally, we investigated the effect of the asymmetric nasal-to-temporal and temporal-to-nasal eye movements on horizontal gaze. Mice have a small frontal region of binocular overlap (about 40 – 50°; Figure 5F) and we wondered to what extent both eyes were aligned with each other during gaze stabilization periods (Figure 5A). Previous work in rats has suggested that alignment can change with head tilt (Wallace et al., 2013). Our data further suggest that alignment can also change during horizontal gaze shifts (in the absence of changes in head tilt; Figure S3C). To quantify the degree of alignment, we computed the difference in horizontal eye velocity between the right and left eyes for periods during which the head was approximately upright (head pitch and roll magnitude < 10°). If both eyes were largely aligned, the distribution of velocity differences would be tightly centered about 0°/s. Instead, we found that distributions for CCW and CW head rotations had a rather wide spread and were shifted and skewed towards the eye that moved temporal-to-nasal (Figure 5G,H; median absolute deviation: 31.42°/s CCW, 29.53°/s CW; median: 14.70°/s CCW, -10.87°/s CW; skewness: -0.53 CCW, 1.09 CW). This suggests that when mice stabilize gaze during head yaw rotations, binocular alignment varies in width, even for horizontal eye movements that were largely conjugate. Thus, in contrast to humans, there seems to be no stable base for continuous stereoscopic depth perception using disparity in freely moving mice.

### Both Types of Eye-head Coupling are Preserved During Visually-guided Behaviors

All of the above measurements in unrestrained mice were made from mice that freely explored a circular or rectangular environment. We wondered whether the observed gaze pattern was preserved when mice interacted with behaviorally salient sensory stimuli or when they engaged in visually-guided behaviors. In humans and also other animals, the presence of relevant stimuli can alter gaze shift patterns, for example during foraging (Land, 2019). To test if this was also true in mice, we performed two different experiments.

In the first experiment, a male mouse with head-mounted camera system initially explored an empty environment. A second male mouse was then placed in the same environment and social interactions were monitored, allowing us to compare gaze shift patterns before and while the other mouse was present in the environment (Video S3). There were no discernible differences in any type of eye-head coupling between the two conditions (Figure S4). However, while social interactions may depend on visual input, particularly during approach (Strasser and Dixon, 1986, Pellis et al., 1996), a wide range of additional, non-visual inputs might be used during social behaviors (Chen and Hong, 2018). For example, it is clear from the supplementary video (Video S3) that the mouse closed its eyes for much of the time that his head was close to the intruder mouse even when the interaction involved rapid chasing around the box.

We therefore designed a visually-guided task that resembles some aspects of typical mouse behavior - detection, approach, and tracking - but in contrast to natural behaviors relied exclusively on vision. The visual target was a black rectangle appearing on an LCD display (Figure 6A, Video S4). The mouse could only solve the task by using the visual information on the display, allowing us to isolate the effect of salient visual input. The rectangle randomly appeared at one of two locations and once touched by the mouse moved randomly to the left or right. The mouse had to press the rectangle again within 2 seconds after the rectangle stopped moving to get a drop of soy milk reward at the other end of the box. Touches of the mouse were detected with an infrared (IR) touchscreen mounted on top of the display as previously described (Bussey et al., 1997, Mar et al., 2013, Horner et al., 2013).

**Figure 6:**
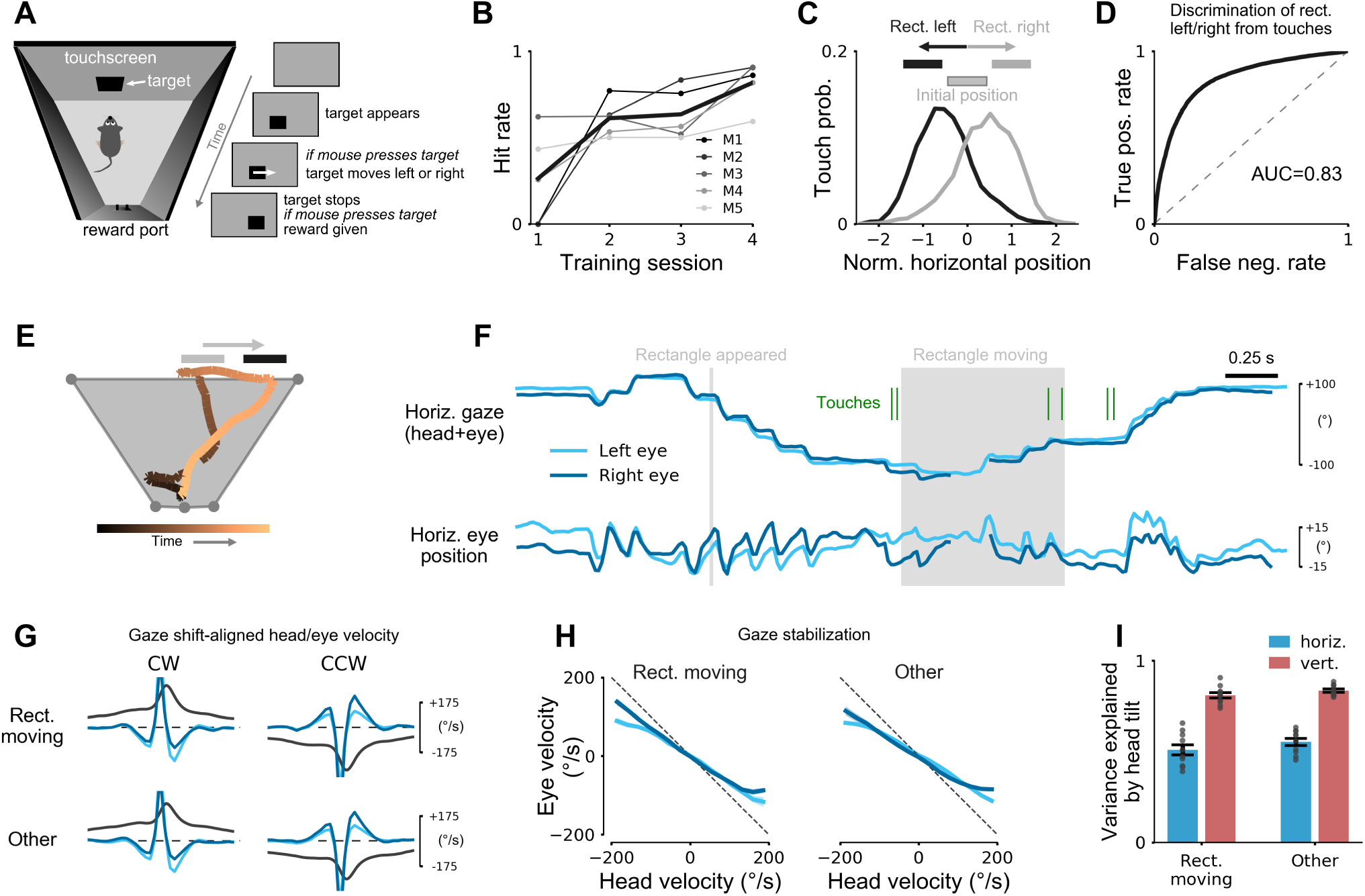
Both Types of Eye-head Coupling are Preserved During Visually-guided Behaviors. (A) Visually-guided tracking task. Mice pressed a black rectangle that appeared on an IR touchscreen. The rectangle then moved randomly for different distances to the left or right and the mouse had to press the rectangle again once the rectangle stopped moving to get a reward at the other end of the box. Mice were first pretrained to press rectangles appearing on the screen with a single touch, and then to press the rectangle for a second time after it had shifted to a new position (B) Learning of the final version of the task in which the initial and final position of the rectangle were non-overlapping. Data shows average hit rates for 5 mice (thin lines) and average hit rate (fraction correct) across mice (thick black line). 211.1 ± 198.9 trials per session. (C) Distribution of touchscreen touches for rectangle moving left (black line) or right (gray line). Touch positions are normalized by rectangle position and width. Extent of rectangles shown above. Data for 4221 trials from 5 mice. (D) Receiver operating characteristics curve for discrimination of left/right rectangle movement based on touchscreen touches for the data shown in C. Area under curve was 0.83. (E) Example trial of mouse performing the task. Overhead view of head position with color indicating trial time as in color bar below. Gray and black rectangles show initial and final rectangle positions, respectively. (F) Gaze (top) and eye (bottom) positions for the trial in E. Rectangle appearance and movement period marked by gray areas. Green lines indicate time points when the mouse is touching the rectangle. (G) Eye-head coupling during gaze shifts was preserved during rectangle tracking compared to a baseline condition (“Other”; without visual stimulus). (H) Relation between head and eye velocity during gaze stabilization periods. (I) Cross-validated explained variance of models trained on head pitch/roll for the baseline condition (“Other”, without visual stimulus). See also Figure S4D,E and Video S4.

Food-restricted mice learned within 3 – 5 days (Figure 6B) to reliably track the object moving left or right (Figure 6C,D). Simultaneous tracking of head and eye movement during behavior enabled us to determine whether eye-head coupling changed when animals tracked a relevant visual object (Figure 6E,F). We observed that the pattern of gaze shifts was similar during the visual tracking task compared to a baseline condition without visual stimulus (Figure 6F–H; permutation tests, all p-values > 0.13; see STAR Methods). There was no significant change in gaze shift frequency between the two conditions (3.0 ± 0.9 gaze shifts per second “Rectangle moving” vs. 3.1 ± 1.1 gaze shifts per second “Other”; Wilcoxon rank-sum test, *p* = 0.87). Moreover, the coupling of eye position to changes in head tilt was preserved; computational models trained on the baseline condition predicted equally well horizontal and vertical eye position from head tilt during stimulus tracking (Figure 6I; *p* = 0.32 horizontal, *p* = 0.23 vertical, Wilcoxon signed-rank test, *n* = 10 (5 mice, left and right eye)).

In sum, both types of eye-head coupling appeared to be maintained when mice were engaged in visually-guided behaviors. How the mice moved their heads during the visual tracking task, however, became highly structured and differed substantially from the patterns observed during free exploration (Figure S4D,E).

### Saccades in Head-restrained Mice Occur During Head Rotation Attempts

Our data suggest that the main role of saccades in mice is to shift gaze during head rotations. We therefore reasoned that saccades observed in head-restrained mice might be linked to the attempt of mice to rotate their heads. To directly test this, we designed an experiment that allowed us to measure eye movements in head-restrained mice along with head rotation attempts without actual motion of the head (Figure 7A). This excluded movement-related signals, such as visual or vestibular input, that could themselves drive eye movements (Van Alphen et al., 2001, Stahl, 2004).

**Figure 7:**
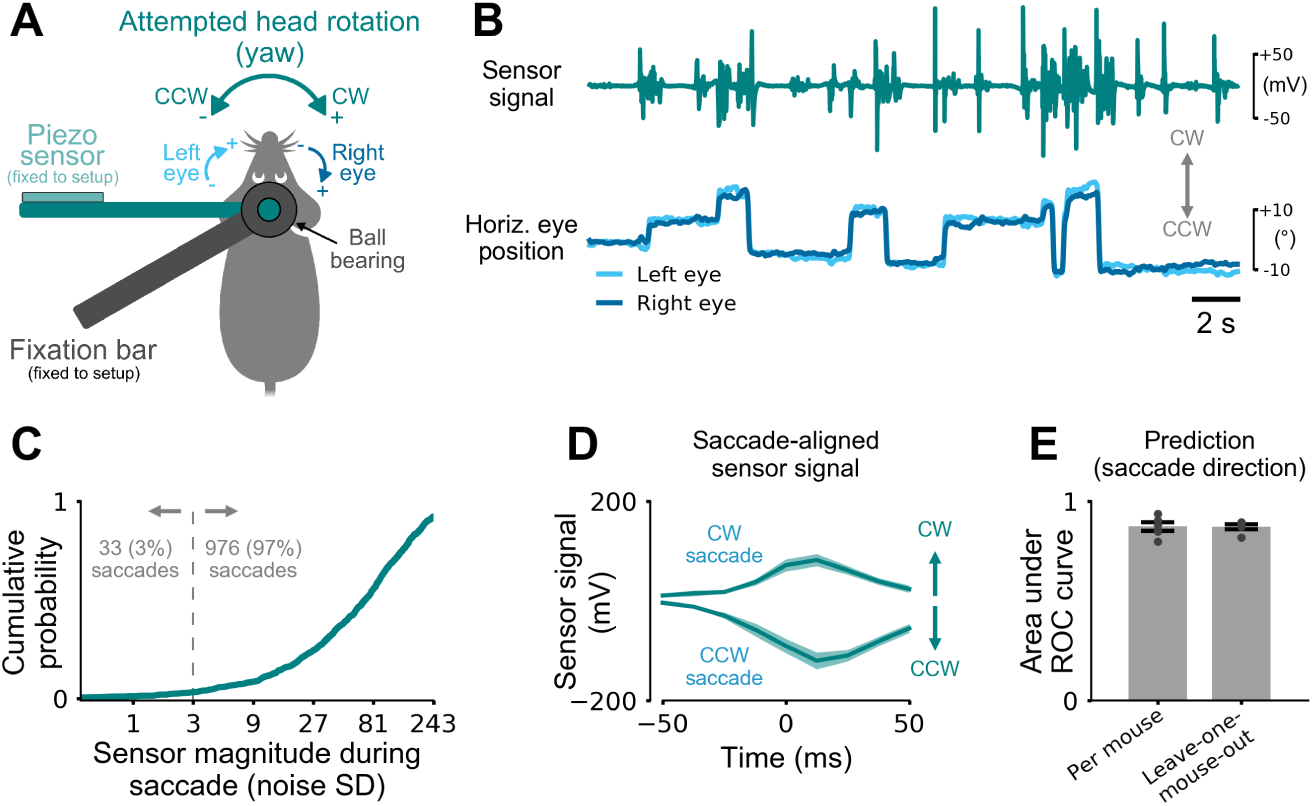
Saccades in Head-restrained Mice Occur During Head Rotation Attempts. (A) Measurement of attempted head rotations in a head-fixed mouse. A fixation bar (dark gray) is attached to the animal’s head post via a ball bearing. A second bar connected to animal’s head post is free to rotate about the yaw axis. The end of the bar is attached to a non-elastic piezoelectric sensor that measures changes in exerted head motion (in the absence of actual head rotation). The animal’s body was restrained by two plastic side plates and a cover above the animal (not shown). (B) Sensor output signal (top) and simultaneously measured horizontal eye positions of both eyes (bottom). Gray arrows indicate CW and CCW directions of sensor signal (head) and eye movements. (C) Sensor signal magnitude during saccades normalized by the standard deviation (SD) of the sensor noise (measured without mouse attached). For 97% of all 1009 saccades in five mice the sensor magnitude was larger than 3 noise standard deviations (dashed gray line). (D) Saccade-aligned sensor trace for CW and CCW saccade directions. Average sensor deflections were in the same direction as the saccades. Mean ± SEM. Same data as in C. (E) Cross-validated prediction performance of saccade directions based on sensor data. Predictions were performed by training a linear classifier using the sensor signals around the saccades (–50 ms to +50 ms, 9 equally-space time points). “Per mouse”: 5-fold cross-validation for each mouse separately. “Leave-one-mouse-out”: saccade direction of a given mouse is predicted using a classifier trained on the data of the other mice. Same data as in C. See also Video S5.

Mice made spontaneous saccades with amplitudes comparable to previous studies (Meyer et al., 2018, Samonds et al., 2018) (Figure 7B; Video S5; horizontal saccade size 12.15 ± 5.11°; 1009 saccades in 5 mice; all values for left eye). During 97 % of all saccades, the sensor used to measure attempted head motion showed fluctuations clearly discernible from baseline (Figure 7C; sensor signal magnitude ≥ 3 sensor noise standard deviations). This indicated that head-restrained mice indeed attempted to rotate their head about the yaw axis during the saccades. The reverse conclusion, however, was not true: not every head movement attempt resulted in a saccade (Figure 7B; Video S5).

We wondered whether these head rotation attempts reflected the same “saccade and fixate” eye-head coupling observed in freely moving mice (Figure 4C). If this were true, then head rotation and horizontal saccade directions should be identical (CW or CCW). To test this, we first aligned sensor signals to either CW or CCW saccades (Figure 7D). Consistent with our findings in freely moving mice, average sensor traces indicated that head rotation attempts were in the same direction as the ocular saccade. To test if this was also true for single saccades, we trained a linear classifier to predict horizontal saccade direction from the head sensor signal measured around the time of the saccades (9 time points from -50 ms before until +50 ms after each saccade). We found that, for each mouse, rotational head motion sensor signals were highly predictive of saccade direction (Figure 7D; area under the ROC curve 0.87±0.02, 5-fold cross-validation; 5 mice). We also tested whether these predictions resulted from patterns that were consistent across mice as suggested by the eye-head coupling in freely moving mice. Predicting saccade direction for each mouse using a classifier trained on data from all other mice (“Leave-one-mouse-out”) resulted in the same high prediction performance (Figure 7E; area under ROC curve 0.87± 0.01) indicating that saccades in head-restrained mice occur during head motion patterns that are similar across mice.

Finally, we tested whether head sensor signals were also predictive of the size of the saccades. We used Bayesian linear regression to predict changes in eye position during saccades (i.e. direction and size) from head sensor traces. Predictions were far above chance level (*r*^2^ = 0.33 ± 0.03) even for leave-one-mouse-out cross-validation (*r*^2^ = 0.33 ± 0.05).

In summary, not only were saccades in head-restrained mice linked to head motion attempts but the patterns were also strongly predictive of saccade direction and size. This shows that rotational eye-head coupling is maintained during head-restraint and suggests that saccades in head-restrained animals might not serve active visual exploration independent of the head.

## Discussion

We have shown that eye movements in freely moving mice consist of two dissociable types (Table 1). By simultaneously tracking eye and head movements in freely behaving mice (Meyer et al., 2018), we find that both types are invariably coupled to the head. The first type, “head tilt compensation”, serves to approximately maintain the same visual field relative to the horizontal ground plane, by systematically changing eye position depending on the tilt of the animal’s head. The second type, the “saccade and fixate” gaze pattern, enables gaze stabilization and gaze shift during reorienting head yaw rotations. Both types, linking eye and head movement, are consistent across a wide range of behaviors being maintained unmodified for example during visually-guided behaviors. This link is so strong that it persists despite attempts to frustrate it: saccadic eye movements in head-restrained mice are associated with attempted head rotation, similar to eye movements in freely moving animals being associated with actual head yaw rotation. These results thereby resolve seemingly contradictory findings of conjugate eye movements in head-restrained mice (Sakatani and Isa, 2004, 2007, Samonds et al., 2018) and complex combinations of conjugate and disconjugate eye movements in freely moving mice (Payne and Raymond, 2017, Meyer et al., 2018), by separating eye movements into two types of eye-head coupling each with its own different linkage. In freely-behaving mice, our results now enable decomposition of the complex eye movement patterns into these two distinct and predictable types where the second saccade and fixate type is further decomposed into two different phases: gaze shift (“saccade”) and gaze stabilization (“fixate”). This helps to clarify which aspects of visual behaviors in humans and non-human primates can be studied in the mouse, which has become a prominent model animal in vision in recent years. Our data highlight five major aspects of mouse eye movements relevant for this comparison.

**Table 1:** The two types of eye-head coupling identified in this study in freely moving mice.

| Type | Description | Conjugate eye movements | Function | Gaze stabilizing | Speed |
| --- | --- | --- | --- | --- | --- |
| 1 | Head tilt compensation | No | Stabilize gaze relative to horizontal plane | Yes | Slow (slower than head) |
| 2 | Saccade and fixate | Yes | “Saccade”: eyes shift gaze together with head during head yaw rotation | No | Fast |
| | | | “Fixate”: eyes stabilize gaze between gaze shifts by counter-rotating against head | Yes | Intermediate (≈ head) |

### 1. Head tilt compensation eye movements

We found that the average position of the mouse eye strongly depends on the tilt of the head, consistent with previous work in mice (Oommen and Stahl, 2008, Meyer et al., 2018) and rats (Wallace et al., 2013). A distinct characteristic of this head tilt-related eye-head coupling is that the two eyes are non-conjugate, typically converging or diverging from each other when an animal is pitching or rolling its head, a behavior that is different from humans and non-human primates. As a consequence, the mouse maintains a relatively narrow range of angles between the ground plane and its gaze, much narrower than the range of angles between the ground and the axis of the eye in its head reference frame. This compensatory process has also been observed in head-restrained mice and has been suggested to reflect preferred alignment of specific parts of the visual field with specific locations on the retina (Oommen and Stahl, 2008).

Here we focused on changes in horizontal and vertical eye position, since they are the main determinants of the visual field (Oommen and Stahl, 2008). Future studies could extend the results to measurements of eye torsion, although torsion in mice is difficult to estimate non-invasively with video-based methods with little retinal structure to provide reference (Oommen and Stahl, 2008) (but see Wallace et al. (2013) for eye torsion measurement in rats). For ground-dwelling animals such as the mouse, the horizon typically divides the world into the ground, where for example food and mates are likely to be found, and an upper visual field covering the sky, a region where aerial predators might appear (Yilmaz and Meister, 2013). Indeed, recent evidence suggests that dorsal-ventral shifts in color (Applebury et al., 2000, Szatko et al., 2019) and contrast (Baden et al., 2013) sensitivity fall onto the ground-observing dorsal and sky-observing ventral retina, respectively. This is in agreement with our finding that the head tilt compensation eye system is trying to keep these parts of the retina aligned with the horizontal plane, regardless of other aspects of behavior or the presence of a salient stimulus as in our visual tracking task. At the same time, due to the arrangement of the eyes in the head, the same eye-head coupling may also help to ensure continuous coverage of a large fraction of the animal’s visual field by the two eyes as reported for rats (Wallace et al., 2013).

Finally, rodents do not rely solely on vision for sensory exploration of the immediate environment but also on sniffing and whisking (Welker, 1964), both of which are coordinated with rapid rhythmical head movements (Deschenes et al., 2012). The head might therefore act as a common reference frame to coordinate information from the different senses.

### 2. Saccade and fixate movements

Horizontal gaze involves fixations in which gaze is approximately kept still, interspersed with saccades to rapidly change gaze direction together with the head. This “saccade and fixate” pattern is observed in many vertebrates, including humans, and enables both stable fixation and rapid gaze shifting with minimal retinal blur (Land, 2019). In a classic paper, Walls argued that the origin of eye movements lies in the need to keep an object fixed on the retina, not in the need to scan the surroundings (Walls, 1962). Indeed, we show that a major aspect of mouse eye movements is the stabilization of retinal stimulation during head rotation. Gaze shifting saccades were typically coupled to head rotations and mice made about 1–2 gaze shifts per second, similar to humans (Einhäuser et al., 2009). We considered the possibility that these horizontal gaze movements reflected overt visual attention. The fact that they did not vary in number or property between spontaneous locomotion in an open field and a visual tracking task would seem to mitigate against this hypothesis. Thus, in the mouse, changes in overt visual attention behavior appear to be mediated by changes in head movements directed to the visual stimulus with eye movements that follow.

### 3. Ocular saccades in head-restrained mice

The possibility that saccades observed in head-restrained mice would normally be associated with head movements during natural behaviors has been suggested previously (Sakatani and Isa, 2007). Correlations between neck muscle activity and eye movements have been shown in different animals, including rabbits (Collewijn, 1977), cats (Vidal et al., 1982, Paré and Guitton, 1990), and primates (Lestienne et al., 1984, Freedman, 2008), while the degree to which the coupling is compulsory appears to vary across species (and appears for example stronger in cats than in primates (Guitton, 1992)). Our data suggest that the coupling in mice is very strong: even in head-restrained mice saccades are invariably associated with attempted head motions. These findings have implications for neural recording experiments in head-restrained animals. If eye movements in these preparations are coupled to and proceeded by head movements, it is necessary to take this into account when searching for neural correlates of eye movements. Correlations may in fact be with other aspects of movement, for example neck proprioceptive or muscle activity, or head movement-corollary discharge signals. Experiments demonstrating changes in the timing of neural signals to eye movements might simply reflect shifts in the correlation from eye to other, head-related movements.

### 4. Strength and consistency of eye-head coupling in the mouse across behaviors

We found that eye-head coupling appears to be relatively consistent across behaviors in the mouse. In other species, including cats and humans, the sequence and contributions of head and eye movements during gaze shifts can strongly depend on the task and the nature of the sensory stimulus (Guitton, 1992). We therefore considered the possibility that eye movements in mice are fundamentally different during visually-guided tasks compared to baseline conditions. Previous studies demonstrated that freely moving mice rely on vision during a variety of naturalistic visually-guided tasks, including social behavior (Strasser and Dixon, 1986, Pellis et al., 1996), prey detection and capture (Hoy et al., 2016), and threat detection (Yilmaz and Meister, 2013). We compared eye movements during baseline conditions, during social behavior (which also relies on other sensory modalities (see Chen and Hong (2018) for a recent review), and during a task that could only be solved using vision that captures aspects of visual detection, approach and tracking in naturalistic mouse behaviors. Our results indicate that during these three conditions, both types of eye-head coupling are maintained; further there was no evidence that saccades lead the head during gaze shifts towards a visual target as in humans and non-human primates (Guitton, 1992, Freedman, 2008, Land, 2015, 2019). This suggests that in the mouse these patterns are less flexible and potentially hardwired for a wide range of visual tasks. There may be good reasons why the mouse gaze system shifts both the eyes and head as a general rule. As mentioned above, the mouse already has a large field of view, which is only shifted by a small amount (approx. 5%) by eye movements alone. Moreover, similar to other animals like cats (Guitton et al., 1984, Guitton, 1992, Einhäuser et al., 2009) and marmosets (Mitchell et al., 2014), mice may be able to rely more heavily on head movement to shift gaze, since they can move the head much faster than monkeys and humans with bigger heads that need to overcome much larger inertial forces. Finally, relying as much as possible on head movements as opposed to eye-in-head movements at the behavioral level reduces the computational burden on the brain to compute this early stage egocentric transformation as it seeks to integrate information from the different sensory modalities in the construction of an allocentric representation of the world.

### 5. Asymmetry in horizontal gaze movements between the two eyes

We discovered a substantial asymmetry in horizontal nasal-temporal and temporal-nasal eye movements in freely moving mice. Thus, similar to head-restrained mice, saccades or stabilizing eye movements along the horizontal eye axis occur simultaneously, but with unequal amplitude in the two eyes (Stahl et al., 2006, Sakatani and Isa, 2007, Itokazu et al., 2018). Since conjugate binocular eye movements typically co-occur with head rotation, the asymmetry might be related to a selective bias for processing visual information in the eye that is on the side of the animal’s heading direction (for example causing improved compensation for head rotation in the left eye compared to the right eye during leftward turns) (Maruta et al., 2006). In any case, without closely yoked eyes, it is not clear how the mouse, a lateral-eyed animal with a narrow binocular field, uses the two eyes to measure distance by disparity during self-motion. Even in the absence of horizontal head and eye movements and the asymmetry noted above, changes in the position of the two eyes as a consequence of head tilt could potentially perturb ocular alignment critical for binocular depth perception (Wallace et al., 2013). Despite this, there is evidence for neural representations of binocular disparities in mouse visual cortex (Scholl et al., 2013, La Chioma et al., 2019). Future experiments could investigate the link between the neural representations of binocular disparity, binocular gaze and visual behaviors in freely moving mice.

### Brain mechanisms

The decomposition of eye and head movements into distinct and independent types has important implications for studying the underlying neural mechanisms. It legitimates the mouse as a useful model to study gaze stabilization and shifting, a prominent feature of visual orienting behavior in humans and other primates. Studying the neural signals during natural visual orienting behavior will require carefully designed experiments to disentangle eye- (Wang et al., 2015, Itokazu et al., 2018) and head motion-related (Wilson et al., 2018) components, and to understand the integration with visual, motor, vestibular, and proprioceptive signals (Chaplin and Margrie, 2019). For example, head tilt stabilization has been reported in the absence of visual input (darkness) (Andreescu et al., 2005, Oommen and Stahl, 2008, Meyer et al., 2018) suggesting that concomitant changes in eye position are driven by vestibular rather than visual input (Andreescu et al., 2005, Oommen and Stahl, 2008). Further, eye movements that stabilize horizontal gaze by moving in the opposite direction to head movement are consistent with the expected effect of the angular vestibulo-ocular reflex. Finally, the superior colliculus, a key structure for controlling head and eye movements in non-human primates (Freedman et al., 1996), likely plays a major role in controlling these movements in rodents (Wang et al., 2015, Wilson et al., 2018, Masullo et al., 2019). Advanced techniques for detailed tracking of head and eye movement (Meyer et al., 2018, Voigts and Harnett, 2019) and virtual reality for visual stimulus control in freely behaving mice (Stowers et al., 2017, Del Grosso et al., 2017), can now be combined with powerful tools to measure and manipulate neural activity (Luo et al., 2008). This provides a unique opportunity to establish the neural circuits that underlie the different types of eye-head coupling.

## Acknowledgments

We thank Maria Chait for support with the human eye tracking experiments; Ben Phillips for advice regarding training of mice on the visual object tracking task; Stephen Burton and Alex Armstrong for their support with the animal work; the Champalimaud Foundation Hardware Platform for providing custom IMU sensor boards; and Bernhard Englitz and Paul Bays for their comments on the manuscript. The authors are also grateful to Jennifer Linden, Maneesh Sahani, Catherine Perrodin, Tim Bussey and Trevor Robbins for helpful discussions and comments during different stages of this work. This work was supported by the Radboud Excellence Initiative (A.F.M), the Wellcome Trust (090843/C/09/Z, 090843/D/09/Z, and 100154/Z/12/A; J.O.K), the Gatsby Charitable Foundation (GAT3212 and GAT3531; J.O.K.), and the UCL Excellence Fellowship (J.P.). J.O.K. is a Wellcome Trust Principal Research Fellow (203020/Z/16/Z). J.P. is a Wellcome Trust and Royal Society Sir Henry Dale Fellow (211258/Z/18/Z).

## Author contributions

Conceptualization, Investigation, and Writing – Original Draft, A.F.M. and J.P.; Methodology, Software, and Data Curation, A.F.M.; Formal Analysis and Visualization, A.F.M., J.P.; Supervision, Resources, and Writing – Review & Editing, A.F.M., J.O.K. and J.P.; Funding Acquisition, A.F.M., J.O.K. and J.P.

## Declaration of Interests

The authors declare no competing interests.

## STAR Methods

### Lead Contact and Materials Availability

Further information and requests for resources and reagents should be directed to and will be fulfilled by the corresponding authors Arne F. Meyer and Jasper Poort.

### Experimental Model and Subject Details

Experiments were performed on five male C57Bl/6J mice (Charles River). After surgical implantation (see “Surgical procedures”), mice were individually housed on a 12-h reversed light-dark cycle (lights off at 12.00 noon).

For the behavioral object tracking experiments, mice had free access to water, but were food deprived to maintain at least 85 percent of their free-feeding body weight (typically 2-3 g of standard food pellets per animal per day). During the other experiments, water and food were available ad libitum. All experiments were performed in healthy mice that had not been used for any previous procedures. All experimental procedures were carried out in accordance with a UK Home Office Project Licence approved under the United Kingdom Animals (Scientific Procedures) Act of 1986.

We also collected eye and head tracking data in 5 human subjects (2 females and 3 males, age 26-51). All gave written consent and the study was approved by the local ethics committee of the Department of Psychology at the University of Cambridge.

### Method Details

#### Surgical procedures

Mice aged 44–49 days were anaesthetized with 1-2% isoflurane in oxygen and injected with analgesia (Carprofen, 5 mg/kg IP). Ophthalmic ointment (Alcon, UK) was applied to the eyes and sterile saline (0.1 ml) injected subcutaneously as needed to maintain hydration. A circular piece of scalp was removed and the underlying skull was cleaned and dried. A custom machined aluminum head-plate was cemented onto the skull using dental adhesive (Superbond C&B, Sun Medical, Japan). Three miniature female connectors (853-87-008-10-001101, Preci-Dip, Switzerland) were fixed to the implant with dental adhesive to enable stable connection of two cameras and an inertial measurement unit (IMU) sensor during experimental sessions. The positions and angles of the two eye tracking cameras and IR mirrors were adjusted using a stereotaxic instrument (Model 963, Kopf Instruments, USA) to align the view of the camera with the eye axis. Additionally, the pitch and roll axes of the IMU sensor were aligned to coincide with the plane spanned by the horizontal eye axes. Mice were allowed to recover from surgery for at least five days and handled before the experiments began.

#### Eye and head tracking in mice

The custom head-mounted eye and head tracking system has been described in detail previously (Meyer et al., 2018). Briefly, we used commercially available camera modules (1937, Adafruit, USA; infrared filter removed), one for each eye. Each camera was inserted into a custom 3D printed camera holder that contained a 21G cannula Coopers Needle Works, UK) to position and hold a 7 mm square IR mirror (Calflex-X NIR-Blocking Filter, Optics Balzers, Germany) and two IR LEDs (VSMB2943GX01, Vishay, USA) to illuminate the camera’s field of view. The camera holder was attached to the connectors on the animal’s head-plate using a miniature connector (852-10-008-10-001101, Preci-Dip, Switzerland). The mirror position was adjusted during surgery (see “Surgical procedures”) and fixed permanently using a thin layer of strong epoxy resin (Araldite Rapid, Araldite, UK) after verifying correct positioning in the head-restrained awake mouse. Camera data were acquired using single-board computers (Raspberry Pi 3 model B, Raspberry Pi Foundation, UK), one for each eye camera, and controlled using a custom plugin (Meyer et al., 2018) for the open-ephys recording system (http://www.open-ephys.org) (Siegle et al., 2017). For all recordings, camera images were 640 x 480 pixels per frame at 60 Hz.

Head rotation and head tilt (pitch and roll) were measured using a calibrated IMU sensor (MPU-9250, InvenSense, USA) mounted onto a custom miniature circuit board with integrated lightweight cable (Champalimaud Foundation Hardware Platform, Lisbon, Portugal). The total weight of the assembly was less than 0.5 g (including suspended part of the cable). Sensor data were acquired at 190 Hz using a microcontroller (Teensy 3.2, PJRC, USA) and recorded along with the camera data using a custom open-ephys plugin (see “Data and Software Availability”). Precise synchronization of IMU and camera data was ensured by hardware trigger signals generated by the microcontroller that were recorded by the open-ephys recording system. The delay of the IMU system was measured by comparing accelerometer signals recorded with the IMU to an analog accelerometer (ADXL335, Analog Devices, USA) recorded directly with the recording system for a number of experiments not included in the analysis. The delay was constant (5 ms) with minimal jitter (<1 ms) and was compensated for prior to the analysis. Head pitch and roll were extracted from accelerometer signals as described previously (Meyer et al., 2018).

All experiments were conducted in a custom double-walled sound-shielded anechoic chamber (Meyer et al., 2018). Animals became accustomed to handling and gentle restraint over two to three days, before they were head-restrained and placed on a custom circular running disk (20 cm diameter, mounted on a rotary encoder). After animals were head-restrained the camera holders and the IMU sensor were connected to the miniature connectors on the animal’s head-plate (see “Surgical procedures”). Power to the infrared light-emitting diodes attached to the camera holders for eye illumination was provided by the IMU sensor.

#### Extraction of pupil positions from camera images

For each eye, we tracked the position of the pupil, defined as its center, together with the nasal and temporal eye corners. Tracking of the eye corners allowed us to automatically align the horizontal eye axis, even in the presence of potential camera image movement (which typically occurs in less than 1% of all frames as shown previously (Meyer et al., 2018)), and to exclude eye blinks. First, about 50–100 randomly selected frames for both eyes (from a total of typically 70000 – 210000 frames) were labeled manually for each recording day. The labeled data were used to train a deep convolutional network via transfer learning using freely available code (https://github.com/AlexEMG/DeepLabCut). The structure and training of the network has been described in detail elsewhere (Mathis et al., 2018). The trained neural network was then used to extract pupil position and eye corners from all video frames independently for each eye. The network predicts the probability that a labeled part, e.g., the pupil center, is in a particular pixel. We used the pixel location with the maximum probability for each labeled part and included only parts into the analysis with probability ≥ 0.99. For our freely moving mouse experiments, this resulted in successful tracking of all labeled parts in 0.95% (1711098/1804120) of frames for the left eye and 96% (1725879/1805361) of frames for the right eye. Video S1 shows examples of tracked pupil center and eye corners for both eyes in a freely moving or head-restrained mouse.

The horizontal eye axis was defined along the line connecting nasal and temporal eye corners; the vertical eye axis was orthogonal to this line. The origin of the eye coordinate system (Figure 2B,D,E) was defined as the mid point between the nasal and temporal eye corners. Pixel values in 2-D video plane were converted to angular eye positions using a model-based approach developed for the C57Bl/6J mouse line used in this study (Sakatani and Isa, 2004).

For the analysis, extracted eye position traces were smoothed using a 3-point Gaussian window with coefficients (0.072, 0.855, 0.072). Eye velocity was computed from the smoothed eye position traces. For analyses that involved comparison of the positions of the two eyes (Figure 1D,H,L and Figure 3C,E), eye position traces were first mapped into the same, uniform time base (sampling frequency 60 Hz) using linear interpolation. The same approach was used to align eye data with head pitch and roll or head angular velocity traces.

For the experiments in freely moving humans, eye position and head motion were recorded using a commercially-available head-mounted eye tracker with integrated IMU sensor (Tobii Pro Glasses 2, Tobii Pro, Sweden). Data were acquired at 100 Hz and analyzed offiine. Before each recording, the eye tracker was calibrated using the supplied calibration routine. Angular horizontal and vertical eye positions were computed from the output of the tracker’s 3D eye model as the angle between the gaze vector and the vector pointing to the front relative to the wearable eye tracker. Care was taken to ensure that the eye tracker was aligned with the frontal plane such that the origin of the angular eye coordinate system (in head tracker coordinates) coincided with the eyes pointing straight ahead (in head coordinates). Eye coordinates were defined analogously to those in mice (Figure 1B): horizontal eye position increases for clockwise eye movements (rotation axis pointing downwards) whereas vertical eye position increases for upward eye movements.

#### Calculation of gaze angle relative to the horizontal plane

Calculation of the angle between the gaze and the horizontal plane required transformations between reference frames. The first was the eye-centered reference frame (“eye in orbit”) represented by horizontal and vertical angular eye coordinates relative to the eye axis, i.e. the origin of the eye coordinate system (see “Extraction of pupil positions from camera images”). The position of the eye axis was defined in a second, head-centered reference frame (“eye in head”). We used an existing geometric model of the eye axes in the head for C57B1/6J mice (Sakatani and Isa, 2007), with the position of the left or right eye axes at ±60° azimuth (relative to midline) and 30° elevation (relative to the animal’s head plate). The third reference frame was the position of the head in the laboratory environment. As the angle between the gaze and the horizontal plane is invariant against rotations about an axis vertical to the horizontal plane, and translations parallel to the horizontal plane, the transformation from a head-centered to a ground-centered reference frame was determined by head pitch and roll (“head in space”; measured using the head-mounted accelerometer).

The transformations between the different reference frames was implemented by multiplication of 3D rotation matrices in the order defined above (“eye in orbit” first, “head in space” last; see also Figure S1A):

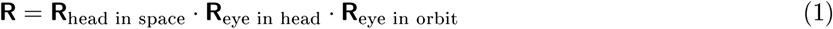

where · denotes matrix multiplication. Each matrix describes elemental rotations about the axes of a Cartesian coordinate system (using the right-hand rule) with axes defined in Figure S1A:

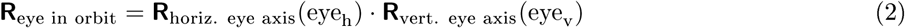

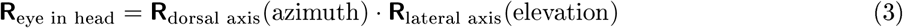

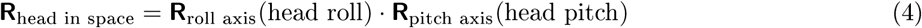

where eye_h_ and eye_v_ denote the current horizontal and vertical angular eye position, azimuth and elevation the orientation of the eye axis in the animal’s head, and head roll and head pitch the current head pitch and roll angles, respectively. Thus, the eye gaze vector, defined as the eye position in space, was computed as

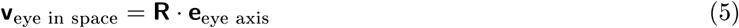

where **e**_eye axis_ = (0, 1, 0)^⊤^ is a unit vector along the eye axis. The gaze angle was defined as the angle between **v**_eye in space_ and the horizontal plane (spanned by the pitch and roll axes in Figure S1A). Similarly, the angle between the eye axis and the horizontal plane was computed by setting eye_h_ = 0° and eye_v_ = 0°.

The resulting transformation was implemented in Python using the numpy and scipy packages (Oliphant, 2007). Video S2 shows an example of the resulting gaze, eye axis, and head tilt vectors together with eye camera frames. Head or eye position angles for which the above configuration could not distinctly be described by the rotation matrices (“Gimbal Lock”) are extremely rare in mice (Wilson et al., 2018) and were excluded from the analysis. We also tested whether the choice of the specific geometric eye axes model affected our results by using a different model (64° azimuth, 22° elevation; (Oommen and Stahl, 2008)). This model gave quantitatively similar results (Figure S1B–D).

#### Prediction of eye position using head tilt

Horizontal and vertical eye positions were predicted from head tilt (pitch and roll) as described in Meyer et al. (2018). Briefly, for each pupil position, the most recent history of head pitch and roll signals within a time window of 100 ms was recast as a single vector containing pitch and roll values. Linear interpolation was used to find pitch/roll at time lags −100, −75, −50, −25, 0 ms. In order to perform nonlinear regression, a Multilayer Perceptron with one hidden layer (100 hidden units with rectified-linear activation functions) was fit to the data. The network was trained using the backpropagation algorithm and weights were optimized using a stochastic gradient-based solver with adaptive momentum estimation via the sklearn Python package (Pedregosa et al., 2011).

The prediction performance of the regression model was evaluated using cross-validation (with *n* = 5 fold). That is, the dataset was divided into 5 parts, model parameters were estimated leaving out one of the parts, and the predictive quality of the model fit was evaluated on the part left out. This procedure was repeated leaving out each of the 5 parts in turn and the prediction accuracy averaged to yield an estimate of the goodness-of-fit of the model. Similarity between predicted and measured eye positions (horizontal or vertical; Figure 3B) was quantified using the coefficient of determination 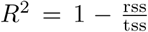 where rss is the residual sum of squares and tss is the total sum of squares. The interocular prediction correlation (Figure 3C) was computed as the correlation coefficient between predictions by independent models for the two eyes. Similarly, the interocular error correlation (Figure 3E) was computed as the correlation coefficient between predictions errors of the two independent models.

#### Extraction of saccades

Saccades were defined as rapid, high-velocity movements occurring in both eyes with magnitude exceeding 350 °/s. Saccade times were extracted from eye velocity traces (see “Extraction of pupil positions from camera images”) by first thresholding eye speed (including eye movement in horizontal and vertical direction). Peak velocity time points were identified by computing local extrema for all time points above the threshold. To avoid double-counting saccades, local extrema were computed in a time window of ±50 ms around the peaks, where 50 ms corresponds to the typical saccade duration in mice (Sakatani and Isa, 2007). Thus if two peaks occurred within 50 ms then only the peak with larger velocity magnitude was classified as a saccade. Relative local extrema were computed using the function “argrelextrema” in the scipy Python package (Oliphant, 2007). Only saccades detected in both eyes, i.e. saccades that were separated in time by less than twice the frame interval of the eye cameras (frame rate 60 Hz), were included in the analysis. Including all saccades detected in each eye regardless of whether a saccade was detected in the other eye revealed the same coupling between the eyes (Figure S2D).

Saccade sizes were computed by first oversampling eye traces around the identified saccades time points (±100 ms) at a time resolution of 5 ms. The start and end points were taken as the points at which the speed of the eye fell below 50 °/s or at ±35 ms if eye speed stayed above this threshold before or after the time point of maximal saccade speed. The saccade sizes reported in Figure 4 were computed as the differences in horizontal eye position at the identified time points after and before each saccade.

#### Quantification of gaze stabilization

The extent to which eye movements stabilized horizontal gaze between gaze shifts was quantified using two different measures. The first measure, mean absolute deviation (MAD), quantified the average deviation from complete compensation (dashed line with slope -1 in Figure 5D,E). The MAD does not make any assumptions about linearity of the relation between head and eye velocity. The second measure, “gain”, was defined as the negative slope of the linear fit between eye and head velocity. A gain of 1 corresponds to full compensation whereas a gain of <1 indicates incomplete compensation suggesting visual slip during gaze stabilization periods. The best linear fit was computed using linear regression on the individual data points (i.e. not the binned data shown in Figure 5D,E).

#### Social interaction experiments

A male mouse with head-mounted cameras and IMU sensor was placed in an empty rectangular environment (length x width x height: 38 cm x 23 cm x 27 cm) and was allowed to freely explore the environment for 10 minutes (“baseline”). A second male mouse without any head-mounted cameras or IMU was then placed in the same environment and social interactions were recorded for approximately 10 minutes (“interaction”). The first mouse had not encountered the second mouse before. Interactions were closely monitored using external video cameras.

During interactions, mice showed a rich behavioral repertoire, including approach, investigation, and attack. Video S3 shows a typical 60 s example segment. Figure S4 shows a summary of horizontal gaze and head tilt-related stabilization quantities extracted from all recordings.

#### Visual motion tracking experiments

We used a custom touch display setup based on an existing design (Horner et al., 2013, Mar et al., 2013). The experiment chamber had a symmetric trapezoidal shape (width: 24 cm on the display side and 6 cm on the side opposite to the display; length: 18 cm; height: 20 cm). The setup was controlled with custom Python software running on a single-board computer (SBC; Raspberry Pi 3B, Raspberry Pi Foundation, UK). Presses of the mouse were detected with an infrared touchscreen mounted onto a 12.1 inch LCD display (NEX121, Nexio, Korea) and read out by the SBC (via serial interface). Nose pokes were detected by the SBC using an IR beam break detector (OPB815WZ, TT Electronics, UK) integrated into a lick spout opposite to the touch display. Soy milk rewards were delivered by opening a pinch valve (161P011, NResearch, USA) connected to the lick spout via silicon tubing (TBGM100, NResearch, USA). Reward delivery was controlled by the SBC via a valve driver (CDS-V01, NResearch, USA).

During training, mice first learned to collect a soy milk reward delivered through the lick spout. No rectangle was shown on the display (gray background). The next day, the delivery of the reward was made contingent on the animal pressing the touchscreen when a full screen black rectangle was shown. Mice had to collect the reward at the other end of the box before the next trial was presented. The size of the rectangle was then gradually shrunk but remained horizontally centered. The vertical position of the rectangle center was 5.8 cm above the ground so mice could easily touch the rectangle with their nose or paws. Once the width of the rectangle was reduced to 5 cm, mice were presented with 3–5 different target positions (3 positions in three mice and 5 positions in two mice, spaced 4 cm apart). As soon as mice were reliably pressing a rectangle appearing at any of the positions (typically after 4–5 days), we trained the mouse on the motion tracking task (Figure 6A). To initiate a trial, mice had to press a rectangle on the screen, which then randomly moved left or right. The horizontal position of the rectangle was computed as

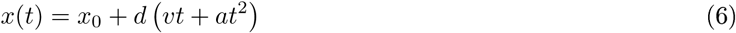

where *x*_0_, *d, v* and *a* denote that rectangle’s initial x-position, moving direction (*d* ∈ {−1, 1}), velocity and acceleration, respectively, and *t* is the time since the rectangle started to move with time steps determined by the frame rate of the LCD display (60 Hz). The rectangle could appear at either *x*_0_ = −2cm or *x*_0_ = +2cm and both *x*_0_ and the moving direction *d* were chosen randomly on every trial such that all four combinations occurred with equal probability. Velocity and acceleration were kept fixed across all experiments (*v* = 2cm/s and *a* = 5cm/s^2^). The mouse had to press the rectangle again at its final position within 2 seconds in order to receive a reward (hit trial). Otherwise, the trial was considered a miss trial. The distance was gradually increased from 4 cm to 6 cm and rectangle width was decreased from 6 cm to 4 cm. Within each experimental block (typically 50 trials), the distance was kept constant. Figure 6B shows hit rate (fraction of hit trials) for blocks without overlap between initial and final rectangle positions (i.e. distance ≥ rectangle width).

The position of the mouse (Figure 6D and Figure S4D,E) was extracted by training a deep convolutional network (Mathis et al., 2018) to detect the locations of the two head-mounted cameras, along with multiple reference points of the setup to automatically map head position onto the geometry of the environment. Head position was defined as the center between the two eye cameras.

To assess the differences between gaze shift patterns in the two experimental conditions (“Rect. moving” and “Other” in Figure 6G) we computed the least absolute deviation (L1 norm) between the gaze shift-aligned head or eye velocity traces for both conditions. To determine the significance of this difference, we used a permutation test. A null distribution was generated by shuffiing “Rect. moving” and “Other” condition labels across all recordings and mice. The permutation procedure was repeated 1000 times, and a P-value was generated by computing the fraction of permutations with least absolute deviations larger than the value computed on the original dataset (*p* = 0.67 left eye CW, *p* = 0.74 right eye CW, *p* = 0.13 head CW, *p* = 0.99 left eye CCW, *p* = 0.5 right eye CCW, *p* = 0.15 head CCW).

#### Measurement of attempted head motion in head-restrained mice

The animal’s body was restrained on a custom 3D printed platform by two plastic side plates covered with soft foam and a cover above the animal. A bar connected to the animal’s head post was free to rotate about the yaw axis. Movement was restricted to yaw rotations by attaching the head post to the inner ring of a ball bearing (608ZZ, NSK, Japan). The other end of the bar was attached to a non-elastic piezoelectric sensor (RS Pro 724-3162, RS Components, UK) that measured changes in exerted head motion relative to the platform (in the absence of actual head rotation). The whole assembly (mouse with head-post and bar) was held by a second bar (“fixation bar” in Figure 7A) that was fixed to the same base as platform and sensor and was attached to the outer ring of the ball bearing. This excluded translational strain from the signal. Thus, when the animal tried to rotate its head about the yaw axis, the bar compressed (CW) or tensioned (CCW) the piezo sensor along the sensor’s axis with effectively zero deflection. This resulted in positive (CW) or negative (CCW) sensor signals. The setup is illustrated in Figure 7A. The output of the piezo sensor was recorded using the analog input of the open-ephys acquisition board (16 bits analog-to-digital converter resolution).

Before each experiment, eye cameras were connected to the animal’s head-plate and the animal was allowed to settle for 5 – 10 minutes before starting data collection. Saccades were extracted as described above (see “Extraction of saccades”). For the analysis, the sensor signal was band-pass filtered between 1 and 50 Hz using a zero-phase second-order Butterworth filter. The minimum detectable head rotation attempt was determined by the noise of the system. To assess this noise, we performed additional measurements on two different days (5 minutes each) without an animal attached. The average noise standard deviation was used to normalize the data in Figure 7C.

Saccade direction (temporal-to-nasal or nasal-to-temporal) was predicted from sensor data using a linear support vector machine (SVM) classifier. For each saccade, the sensor signal in a time window ±50 ms around the saccade (9 time points) was extracted and a SVM was trained to predict saccade direction from the extracted data. Prediction performance was evaluated using 5-fold cross-validation as described above (but using the area under the receiver operating curve as performance metric). This procedure was done separately for data from every mouse (“Per mouse” in Figure 7E) or by training the SVM using all data but leaving out one mouse and predicting saccade directions for the mouse left out (“Leave-one-mouse-out” in Figure 7E). Similarly, prediction of saccade direction and magnitude was done by linear regression between sensor traces and the change in eye horizontal eye position before and after the saccade. The linear regression weights and the offset term were found using Automatic Relevance Determination (MacKay, 1996).

### Quantification and Statistical Analysis

Specifics on the statistical methodologies and software used for various analyses are described in the corresponding sections in Results, figure legends, STAR Methods, and supplemental figures. Statistical test results are described as significant in the text where P <0.05. All tests were performed using the R software package (version 3.6.1) and the scipy Python package (version l.4.l).

### Data and Code Availability

Code for camera and IMU data acquisition and plugins for controlling the camera (https://github.com/arnefmeyer/RPiCameraPlugin) and IMU (https://github.com/arnefmeyer/IMUReaderPlugin) have been made freely available. Instructions for construction of the eye and head tracking system are publicly available, together with code for extraction of head pitch and roll from accelerometer signals (Meyer et al., 2018). Pupil tracking code and example data will be made available upon publication of the manuscript.

## Supplemental Information

**Figure S1:**
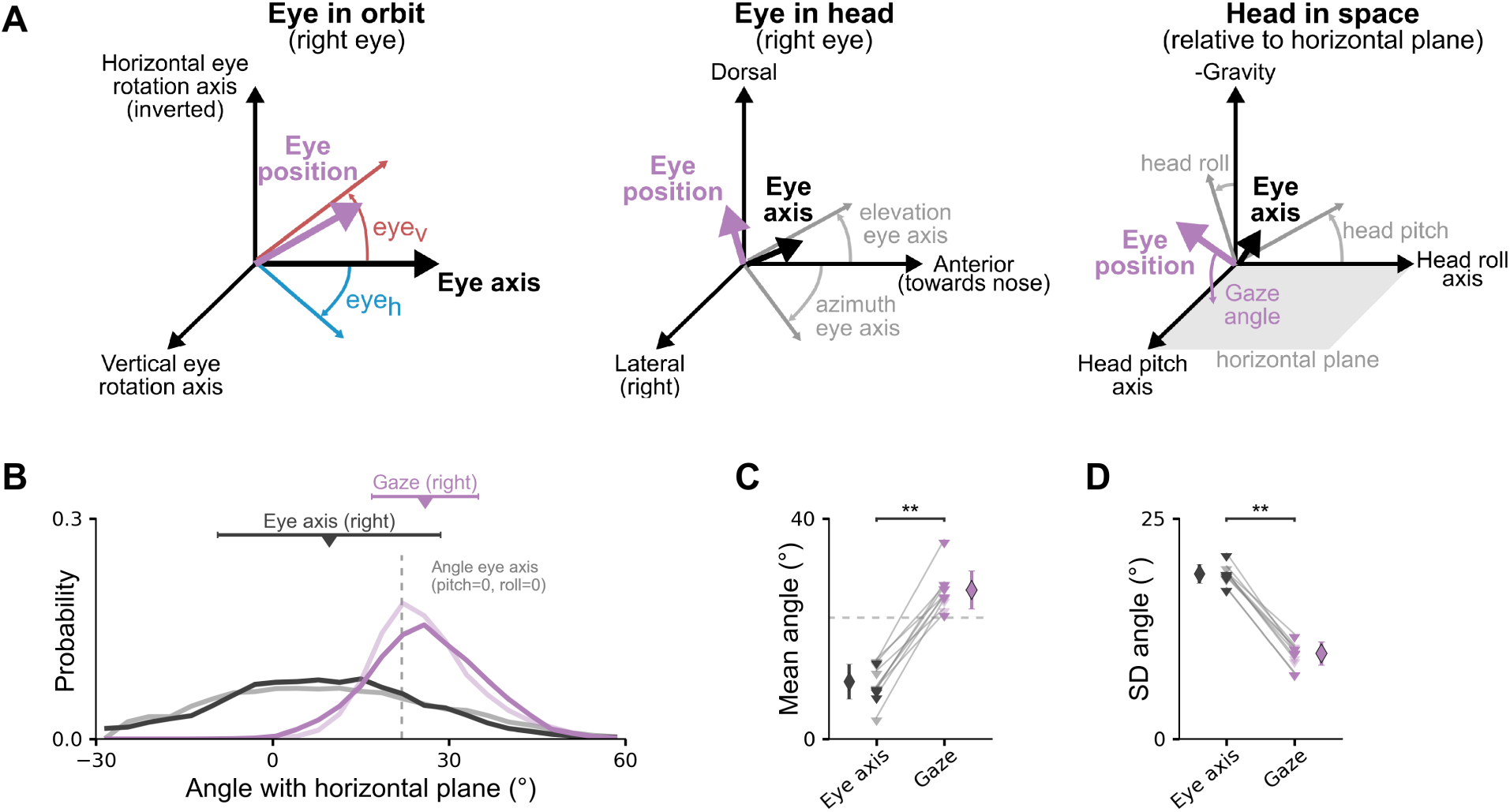
Computation of gaze angle relative to the horizontal plane. Related to Figure 2. (A) Rotations in three different reference frames for computing the gaze angle (the angle between the eye position vector and the horizontal plane in “Head in space”). Rotations were performed in the order from left (“Eye in orbit”) to right (“Head in space”) as described in STAR Methods. Examples show rotation angles for right eye. (B–D) The same as in Figure 2F–H but for the geometric eye axis model described in Oommen and Stahl (2008) (64° azimuth, 22° elevation).

**Figure S2:**
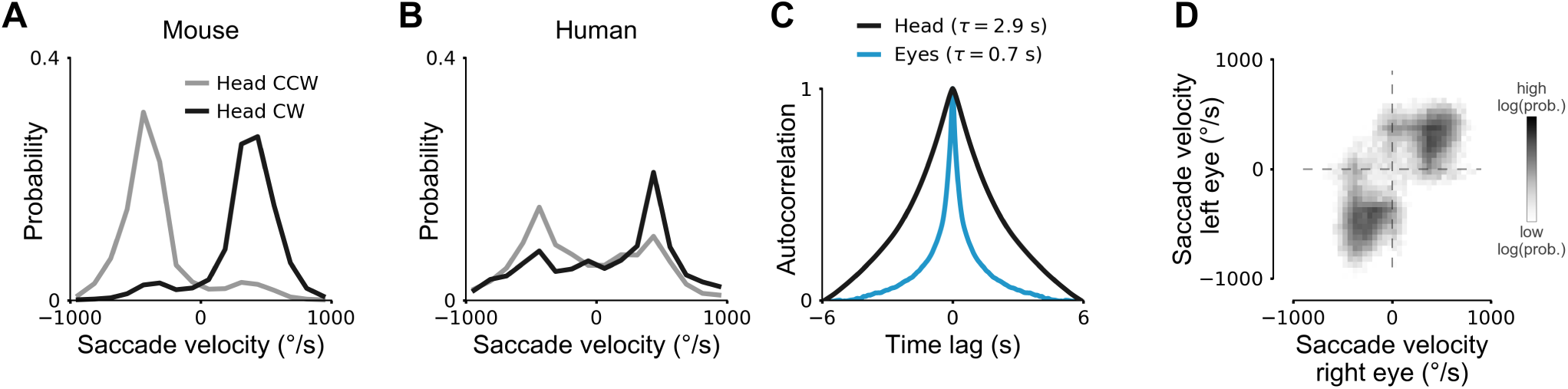
Saccades in freely moving mice and humans. Related to Figure 4. (A) Distribution of saccade velocity for CW and CCW head rotations in freely moving mice. Same data as in Figure 4B–G. (B) The same as in A but for humans walking around the environment. Similar to mice, combined eye-head gaze shifts occur with high probability as indicated by the peaks in the distributions. (C) Autocorrelation function of horizontal eye position (averaged across both eyes) and angular head position (computed by integrating head rotation velocity about the yaw axis). Eye movements have a substantially shorter correlation time constant (*τ* = 0.7 s) than head movements (*τ* = 2.9 s). Time constants were computed by fitting a decaying exponential to the positive lags of the angular horizontal eye or head yaw position correlation functions. (D) The same as in Figure 4B but when including all saccades detected in the left or right eye regardless of whether a saccade was detected in the other eye (see STAR Methods).

**Figure S3:**
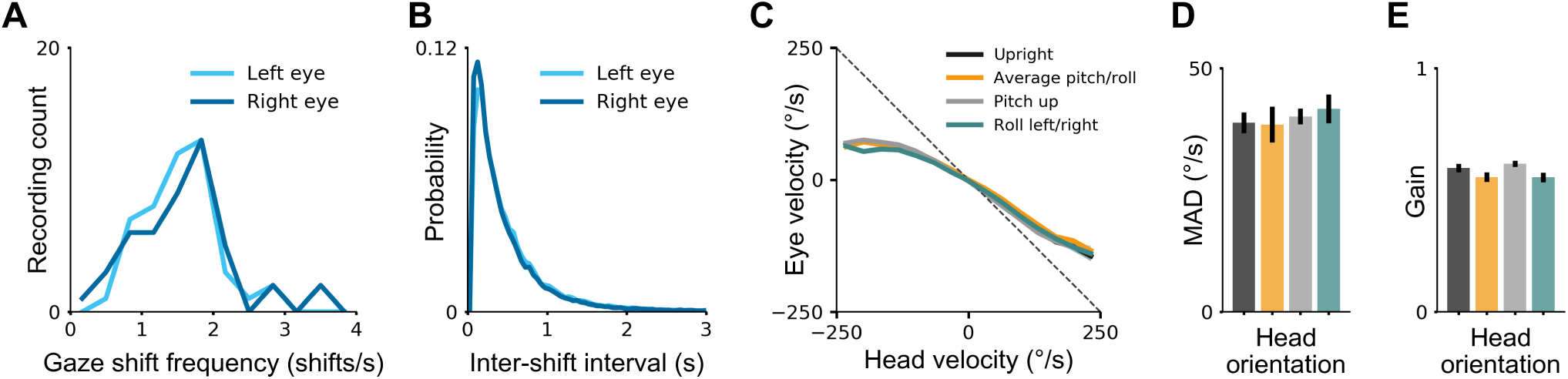
Gaze shift patterns in freely moving mice. Related to Figure 5. (A) Distribution of gaze shift frequency for gaze shifts detected separately for left and right eyes. Data from 47 recordings (each 10 minutes) in 5 mice. (B) Distribution of inter-shift intervals. Peak of distribution at about 150 ms. Same data as in A. (C) Relation between head and eye velocity for different head orientations. “Upright”: head pitch and roll within ±10° relative to vertical (gravity) axis. “Average pitch/roll”: head pitch and roll within ±10° relative to average pitch/roll axis. “Pitch down”: head pitch negative and roll within ±10° relative to vertical (gravity) axis. “Roll left/right”: head roll greater than ±10° away from vertical axis. Saccades were excluded from the analysis. Plots show mean ± SEM for the left eye in 5 mice (typically smaller than line width). (D) Mean absolute deviation (MAD) for left and right eyes in 5 mice. Plots show mean ± SEM. Same color scheme as in C. Differences were small and statistically not significant (Wilcoxon rank-sum test with *α* = 0.05; Bonferroni correction) (E) The same as in D but for horizontal gaze stabilization “gain”. All differences were statistically not significant (Wilcoxon rank-sum test with *α* = 0.05; Bonferroni correction)

**Figure S4:**
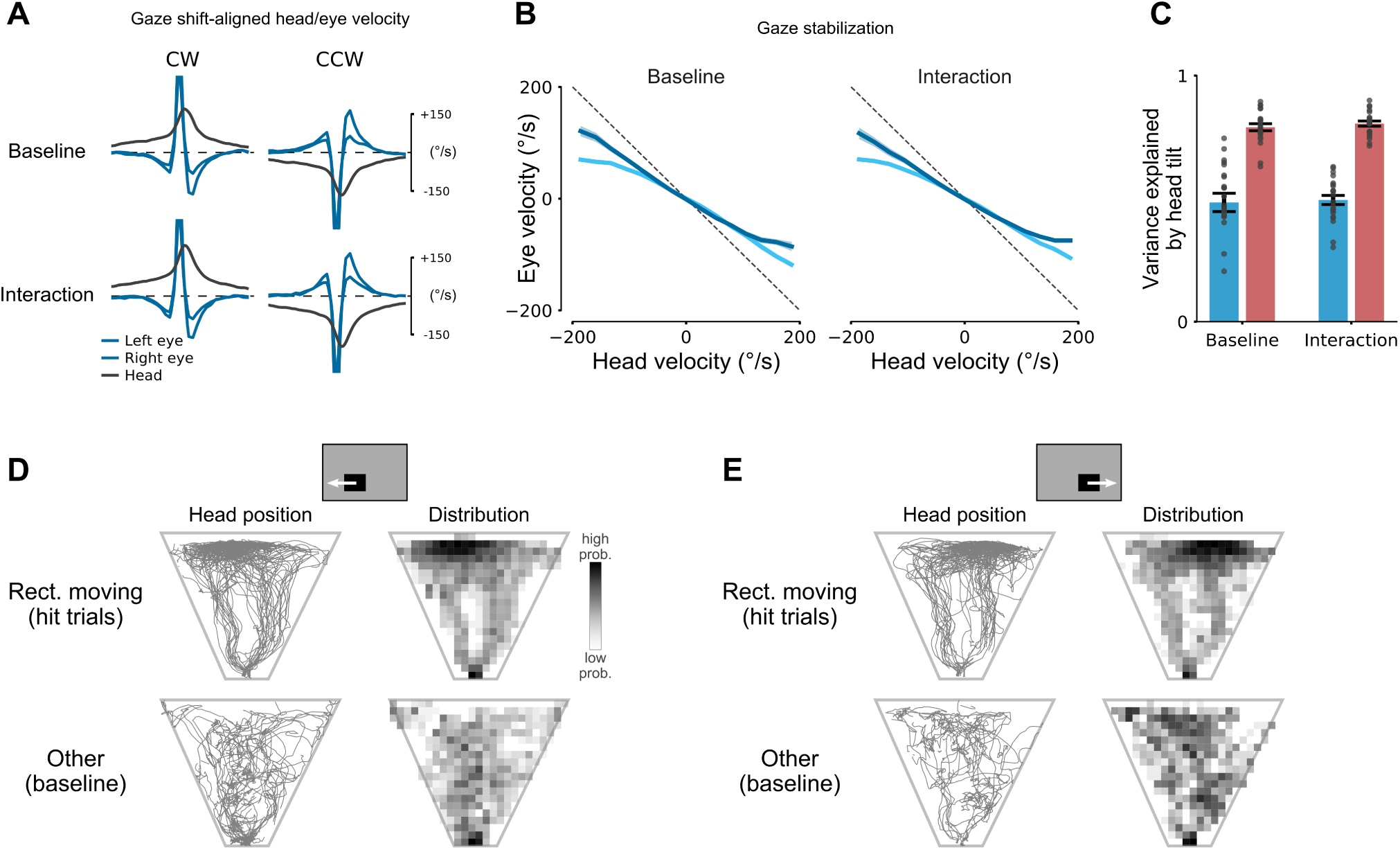
Eye-head coupling during visually-guided behaviors. Related to Figure 6. (A) Gaze shift-aligned head and eye velocity traces measured during social interaction (“Interaction”) and a baseline condition (“Baseline”, without the other mouse in the rectangular environment) recorded before the interaction. 11 paired interaction/baseline recordings in 5 mice (duration interaction 597 ± 11 s, duration baseline 601 ± 0 s). Same analysis as in Figure 5C. (B) Relation between head and eye velocity during gaze stabilization periods. Same data as in A. (C) Cross-validated explained variance of models trained on head pitch/roll for the baseline condition. There were no significant differences in explained variance in horizontal or vertical eye dimension (*p* = 0.54 horizontal, *p* = 0.39 vertical; Wilcoxon sign-rank test). Same data as in A. (D) Tracking of head position during the visually-guided tracking task (Figure 6). Tracked head position (thin gray lines, left) and distribution of head position (log-scaled spatial distribution, right) for an example mouse performing the rectangle tracking task (“Rect. moving”, top) or during free exploration of the same experimental setup without stimulus shown on the display (“Other”, bottom). Top illustration shows initial rectangle position and rectangle movement direction for 63 hit trials (“Rect. moving” condition). For each trial, head position is shown starting 2 seconds before the initial touch of the rectangle until the first touch after the rectangle reached its final position. (E) The same as in D but for a different initial rectangle position and movement direction (53 trials).

## Supplemental Videos

Video S1: **Eye Movements in Freely Moving and Head-Restrained Mice. Related to Figure 1.** Eye movements measured using two head-mounted cameras for the same mouse when it was freely moving or head-restrained. No stimuli or visual feedback were provided during the head-restrained recording. Dots indicate tracked locations of pupil center (blue), nasal eye corner (orange), and temporal eye corner (green).

Video S2: **Orientations of eye axes and gaze vectors relative to the horizontal plane. Related to Figure 2.** Left: eye images of the two head-mounted cameras (top) and external camera showing the mouse exploring a rectangular environment (bottom). Right: orientation of head (thick green arrow), eye axes (thin white arrows), and gaze axes (purple lines) relative to the horizontal plane. Blue and red arrows indicate horizontal and vertical axes of the eye coordinate system, respectively.

Video S3: **Eye tracking in a mouse during social interaction. Related to Figure 6.** Eye camera images (top) together with images of an external camera (bottom) during mouse social interaction. The male mouse with the head-mounted cameras had not encountered the other male mouse (without cameras) before.

Video S4: **Eye tracking in a mouse performing a visually-guided tracking task. Related to Figure 6.**

Video S5: **Eye movements and attempted head movements in head-restrained mice. Related to Figure 7.** Eye images (top), extracted horizontal eye positions (middle) and simultaneously recorded head movement signal (bottom) for the 50 s segment shown in Figure 7B.

